# Immune cell mitochondrial phenotypes are largely preserved in mitochondrial diseases and do not reflect disease severity

**DOI:** 10.1101/2025.01.18.633635

**Authors:** Cynthia C. Liu, Mangesh Kurade, Anna S. Monzel, Catherine Kelly, Robert-Paul Juster, Caroline Trumpff, Michio Hirano, Martin Picard

## Abstract

**Objective:** To profile immune cell mitochondrial phenotypes in mitochondrial diseases (MitoD) and evaluate how those phenotypes relate to disease manifestations or biomarkers.

**Methods:** We profiled mitochondrial content and oxidative phosphorylation (OxPhos) enzymatic activities in isolated monocytes, lymphocytes, neutrophils, platelets, and mixed peripheral blood mononuclear cells (PBMCs) from 37 individuals with MitoD (m.3243A>G, n=23; single, large-scale mitochondrial DNA (mtDNA) deletions, n=14) and 68 healthy women and men from the Mitochondrial Stress, Brain Imaging, and Epigenetics (MiSBIE) study.

**Results:** We first confirm and quantify robust cell type differences in mitochondrial content, activities of OxPhos complexes I, II, and IV, and the mitochondrial respiratory capacity (MRC) index. In relation to MitoD, neither mitochondrial content nor OxPhos capacity were consistently affected, other than a mild monocyte-specific reduction in complex I (partially mtDNA encoded) relative to complex II (entirely nDNA encoded), consistent with the mtDNA defects examined. Relative to the large differences in cell type-specific mitochondrial phenotypes, differences in MitoD relative to controls were generally small (<25%) across mitochondrial measures. The MitoD biomarkers GDF15 and FGF21, as well as clinical disease severity measures, were most strongly related to mitochondrial abnormalities in platelets, and most weakly related to mitochondrial OxPhos capacity in lymphocytes, which are known to eliminate mtDNA defects. Finally, comparing PBMCs collected in the morning/fasted state to the afternoon/fed state following a stressful experience, we report significant time-dependent changes in mitochondrial biology over the time scale of hours.

**Conclusions:** Overall, these results demonstrate that the dynamic and cell-type specific mitochondrial phenotypes are preserved in mitochondrial diseases and are generally unrelated to the severity of MitoD symptoms.

## INTRODUCTION

There is growing interest in probing mitochondrial health profiles using blood-based bioenergetic measures ^1–3^. Mitochondrial oxidative phosphorylation (OxPhos) capacity deficits and changes in blood immune cells have been identified in neurodegenerative diseases such as Alzheimer’s disease, metabolic diseases such as diabetes, sepsis, psychiatric disorders such as major depressive disorder ^4–6^, and is also related to chronic psychosocial stress ^7^. Whether immune cell OxPhos enzyme activities reflect whole body OxPhos activity is unclear. Existing studies show inconsistent results regarding whether immune and muscle mitochondrial OxPhos capacity are correlated ^8, 9^. Moreover, low coherence in mitochondrial gene expression has been found between different organs and tissues within the human body ^10^, calling into question the reliability of using immune system mitochondria as a proxy for the rest of the body. Thus, it is important to understand if immune cell mitochondrial phenotyping can reflect an individual’s mitochondrial OxPhos capacity, including those with mitochondrial diseases (MitoD).

Current work using blood-based measures of OxPhos complex enzyme activities in mitochondrial disease has been limited. Some studies report reduced OxPhos complex activities in peripheral blood mononuclear cells (PBMCs), platelets, and lymphocytes in patients with various mitochondrial diseases ^11–15^, whereas others report no significant differences ^16^ between MitoD patients and healthy controls. In general, assessments of immune cell OxPhos complex activities offer poor sensitivity and specificity for MitoD^17^. This is likely due to the elimination of mitochondrial DNA (mtDNA) mutations within the immune lineage ^18–21^, in contrast to postmitotic tissues, where mtDNA mutation load increases over time ^22^.

Additionally, how OxPhos profiles in specific immune cell subtypes are affected in MitoD is not well understood. Immune cell subtypes possess distinct OxPhos profiles ^23–27^, but how these profiles are affected in MitoD has yet to be investigated. Purifying selection of MitoD mtDNA mutations also occurs in immune cell types, decreasing mutation load as these cells mature. This is particularly evident in lymphoid cells (as opposed to myeloid lineage cells) in both human patients and mouse models with the m.3243A>G pathogenic variant ^18–21^.

Consequently, immune cells OxPhos enzyme activities may not be affected as severely as in somatic tissues, and OxPhos in immune cell subtypes may be differentially affected by MitoD-causing pathogenic variants. MitoD may also cause changes in relative immune cell subtypes abundance ^28^, further complicating the understanding of the immune OxPhos profile in MitoD, and calling for a better understanding of OxPhos activities across different immune cell types. Besides OxPhos capacity, findings around mtDNA copy number (mtDNAcn) changes in MitoD are also mixed ^17, 21, 29^. Cell-type-specific immune studies also show that mtDNAcn is also tissue and immune cell-type specific ^24, 30, 31^.

Finally, the timing for mitochondrial biology assessment may also be important to understand how immune cells are affected in MitoD. The immune system exhibits diurnal variation in the abundance of circulating cell types ^32–34^, and there may be natural within-person variation in immune cell subtype abundance and mitochondrial enzyme activity (for example, from week-to-week ^24^). Whether time-dependent changes in immune mitochondrial biology are altered in MitoD has not previously been examined.

In the present study, we profiled OxPhos enzyme activities (complexes I, II and IV) together with the mitochondrial content markers citrate synthase activity and mtDNAcn, across isolated monocytes, lymphocytes, neutrophils, platelets, and mixed PBMCs in MitoD and age-matched controls. We quantified differences in mitochondrial phenotypes in a cell type specific manner and systematically examined how immune mitochondrial profiles relate to MitoD severity and disease biomarkers. We also explored how PBMCs OxPhos enzyme activities and mitochondrial content change from morning to afternoon, expanding our understanding of immune mitochondrial biology in relation to mitochondrial diseases.

## RESULTS

### Immune cell subtypes have distinct OxPhos capacities and mitochondrial content

To first establish how mitochondrial biology differs between immune cell types, we profiled multiple features of mitochondrial content and OxPhos capacity among monocytes, lymphocytes, neutrophils, and mixed peripheral blood mononuclear cells (PBMCs) from 37 individuals with mitochondrial diseases (two groups – m.3243A>G, n=23; single, and large-scale mtDNA deletions, n=14) and 68 healthy women and men from the Mitochondrial Stress, Brain Imaging, and Epigenetics (MiSBIE) study (aged 18-60 years, 69% women)^35^. In line with our previous findings in a smaller study ^24^, healthy controls showed significant differences in OxPhos enzyme activities (complex I (CI), complex II (CII), complex IV (CIV)), citrate synthase (CS) activity, mtDNAcn, and mitochondrial respiratory capacity (MRC) between immune cell subtypes (**Fig. 1A-F)**. Platelet enzyme activities were also measured, but whereas immune cell activities can be expressed per nucleated cell, platelet enzyme activities are normalized per protein content and therefore not directly comparable to immune cell populations. Results for platelets are included below.

**Figure 1.**
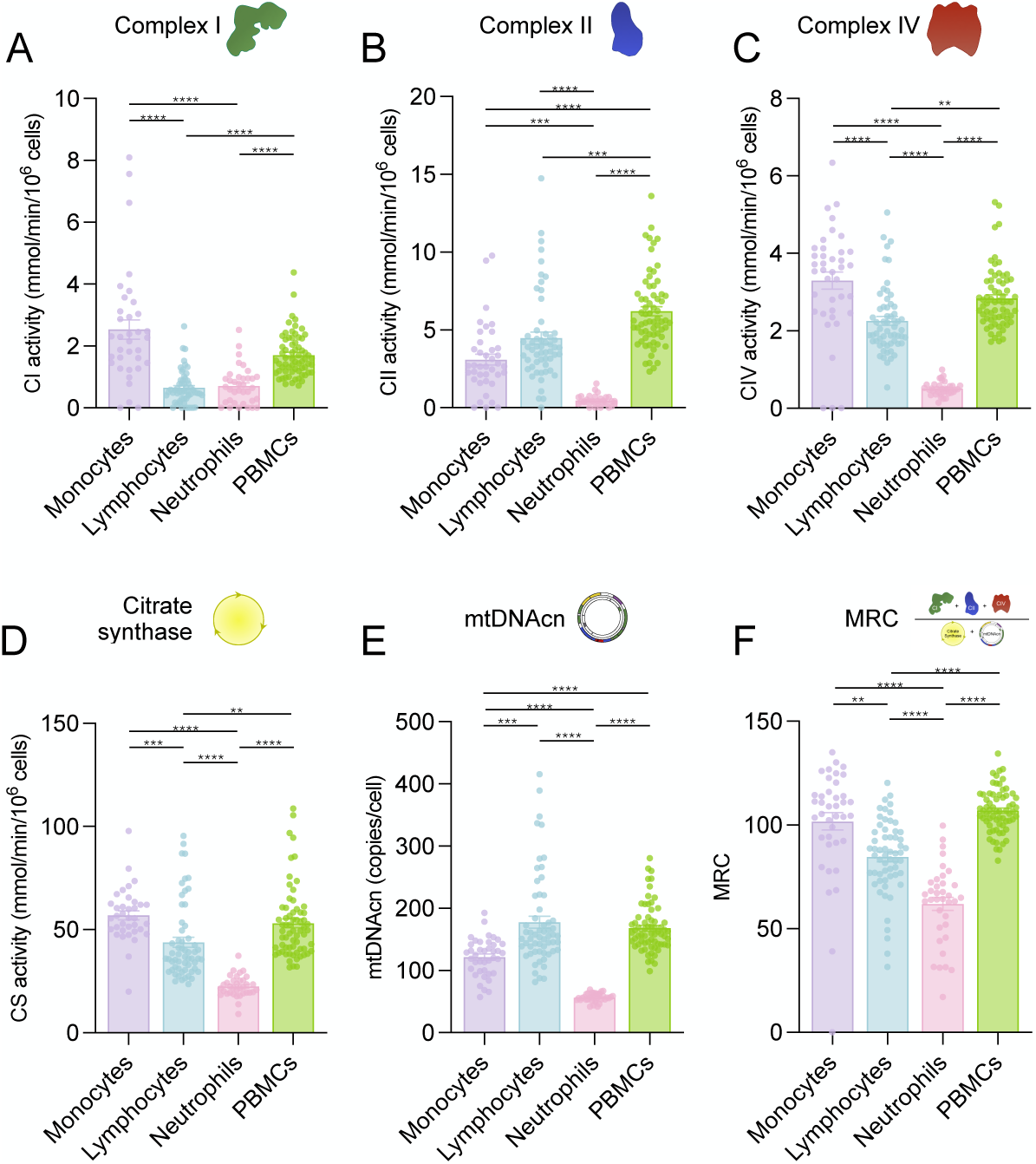
Differences between measures of mitochondrial biology in immune cell subtypes of healthy controls. **(A)** Complex I (CI) activity, **(B)** Complex II activity, **(C)** Complex IV activity, **(D)** Citrate synthase (CS) activity, **(E)** mtDNAcn, and **(F)** Mitochondrial respiratory capacity (MRC) in immune cell types. n=31-66. One-way ANOVA, Kruskal Wallis multiple comparison test. *p<0.05, **p<0.01, ***p<0.001, ****p<0.0001. Individual data points, mean +/- SEM shown.

Of all cell types assessed, in healthy controls, neutrophils had the lowest mitochondrial content (indexed by both CS activity and mtDNAcn) and OxPhos enzymatic activities, consistently less than half that of monocytes and PBMCs, which had high activities across all enzymes. Neutrophils also displayed the lowest mtDNAcn (average 56.6 copies/cell), whereas lymphocytes showed the highest mtDNAcn (average 177.7 copies/cell). The integrative measure of mitochondrial respiratory chain capacity on a per mitochondrion basis, the MRC ^36^, was highest in PBMCs (MRC=107, **Fig. 1F**), likely because of residual platelet contamination that is typical of PBMC preparations ^24, 37^. PBMCs MRC was 5.2% higher than in monocytes (101.7), 26.5% higher than lymphocytes (84.6), and 72.9% higher than neutrophils (MRC=61.9), establishing the spectrum of mitochondrial OxPhos specialization across major immune cell types.

We next assessed whether different mitochondrial features correlated within cell types. As expected, given that the majority of cells in PBMCs are lymphocytes, mitochondrial features tended to correlate well within PBMCs and lymphocytes, (average r’s = 0.59 and 0.56, respectively). Monocytes, neutrophils and platelets did not display strong correlations between mitochondrial features, (average r’s = 0.31, 0.06, 0.06, respectively, **Supp. Fig 1A**), suggesting that these features may be regulated relatively independently of each other in these cell types.

Between cell types of the same individual, we did not find evidence of an association between mitochondrial features, with the exception of CS activity and MRC, which were weakly correlated between neutrophils and PBMCs (r=0.36, p=0.045), and between neutrophils and lymphocytes (r=0.36, p=0.04), respectively (**Supp Fig. 1B, D-G**). However, after correction for multiple comparisons, these correlations were no longer significant (Bonferroni corrected p’s= 0.68 and 0.60, respectively). Thus, individuals with high mitochondrial content or OxPhos activity in one cell type do not necessarily have similarly high mitochondrial features in other cell types, consistent with previous findings ^24^.

For each participant, in addition to the PBMCs collected in the morning/fasted state (PBMCs 1), PBMCs were also collected in the afternoon/fed state following a psychologically stressful laboratory task involving a defense against an alleged shoplifting (PBMCs 2), and the mitochondrial profiling was repeated for samples collected at that time point. As expected, mitochondrial features were significantly correlated between the two PBMC samples (r’s= 0.34 – 0.49, p’s<0.05), for all measures except CS activity (r=0.36, p=0.081) (**Supp Fig. 1B, C**). The overall low correlation between different immune cell types of the same person, but presence of correlation between PBMCs, suggests that mitochondrial respiratory capacity varies by cell type and is not a stable trait of an individual.

### Immune cell types display distinct mitotypes

To further examine the differences in mitochondrial respiratory capacity and content between immune cell types and PBMCs, we created bivariate plots of pairs of mitochondrial features. These plots distinguish between distinct mitochondrial phenotypes, or *mitotypes* ^24^, and allow us to situate different cell types relative to each other in a simple bi-variate mitotype space. These mitotypes are alternatively presented as ratios, where cell types with the same ratios line up along the diagonal, and allow the quantification and statistical comparison of the mitotype profiles of cell types to each other.

We first examined complex IV activity relative to citrate synthase (CIV/CS ratio). Most cell types showed similar CIV/CS ratios, as shown by their alignment along the same y/x line in the CS vs. CIV activity bivariate plot (**Fig. 2A, upper**) and their similar CIV/CS activity ratios (between 0.053-0.064) across most cell types (**Fig. 2B, upper**). Other bivariate mitotype plots revealed significantly larger distinctions between cell types **(Fig. 2, Supp Fig. 2A-K)**. For example, for the CII/CI mitotype reflecting the relative activity of OxPhos complex II (encoded entirely by nDNA genes) relative to complex I (encoded by both the mtDNA and nDNA), lymphocytes displayed low CI activity (0.52 mmol/min/10^6^ cells) but high CII activity (3.87 mmol/min/10^6^ cells), leading to a large CII/CI ratio of 12.4. This positioned lymphocytes in the upper left quadrant, whereas monocytes displayed the opposite pattern, with lower CII activity (2.27 mmol/min/10^6^ cells) relative to a higher CI activity (2.66 mmol/min/10^6^ cells), yielding a ratio of 2.1 and a position in the lower right quadrant of the mitotype plot. Neutrophils were low in both CI activity (0.66 mmol/min/10^6^ cells) and CII activity (0.42 mmol/min/10^6^ cells), with a low average CII/CI ratio of 0.86 and a mitotype position in the lower left quadrant (**Fig. 2A, B, middle**). Immune cell types also differed considerably in their CS/mtDNAcn mitotype, reflecting the mtDNA content per unit of citrate synthase or Krebs cycle-defined mitochondria (**Fig. 2A, B, lower**). Together, these results reinforce the notion that distinct immune cell types have unique mitochondrial properties and respiratory capacities.

**Figure 2.**
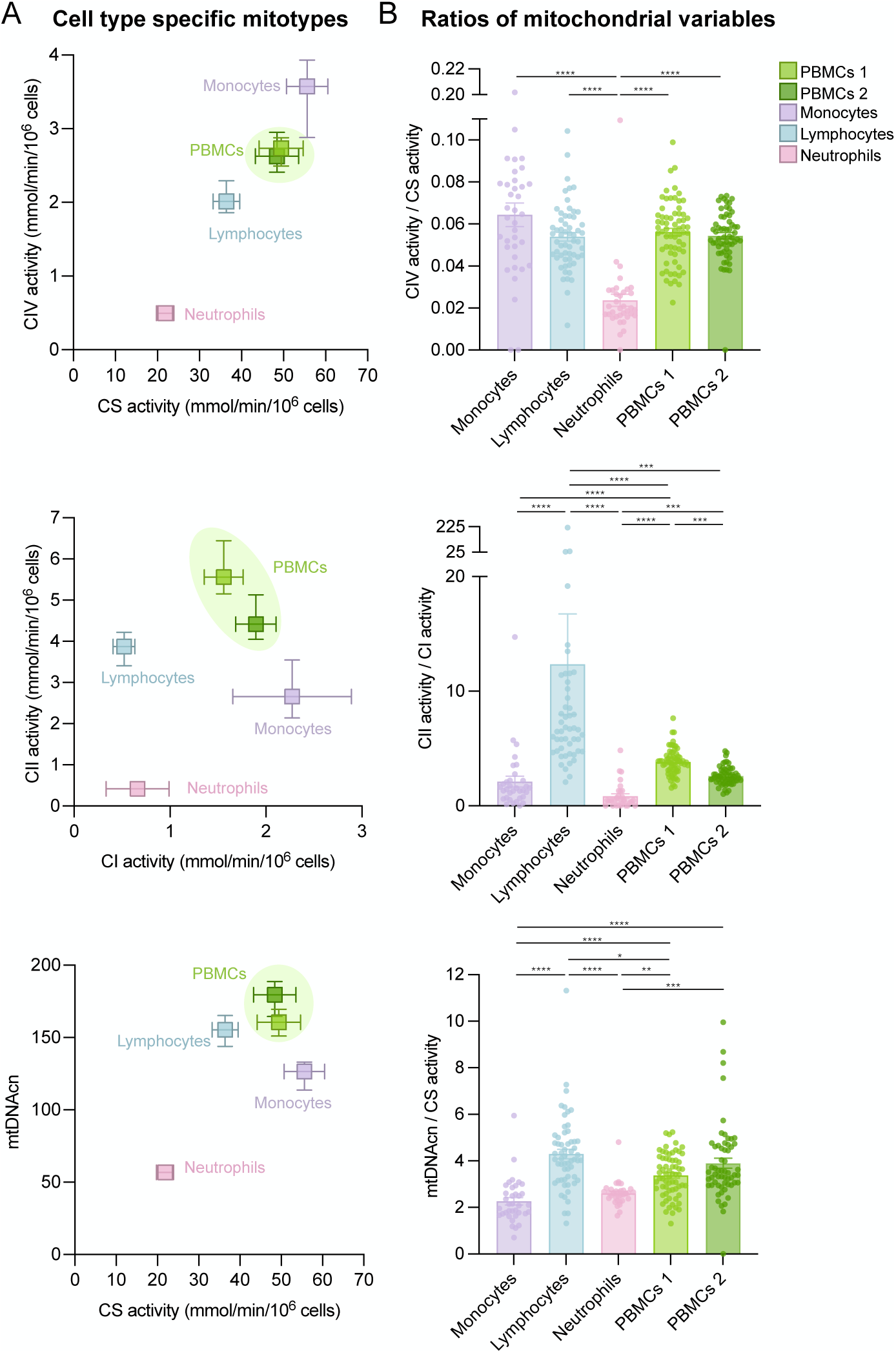
Relationships between different measures of mitochondrial biology by cell type. **(A)** Bivariate plots and **(B)** ratios showing relationship between different pairs of mitochondrial enzyme measures and mtDNAcn in different cell types of healthy controls. PBMCs 1 are PBMCs collected in the morning/fasted state, PBMCs 2 are PBMCs collected in the afternoon/fed state following a psychologically stressful laboratory task. n=31-66. One-way ANOVA, Kruskal Wallis multiple comparison test. *p<0.05, **p<0.01, ***p<0.001, ****p<0.0001. Individual data points, mean +/- SEM shown.

### Mitochondrial OxPhos capacity and content were not affected in mitochondrial disease

We next compared mitochondrial features between MitoD and controls in each of the specific immune cell types. Across all cell types and mitochondrial respiratory capacity measurements, most differences were small (average change 11.7%) and non-significant. The only significant difference between control and MitoD groups was a 32% higher CIV activity in monocytes of MitoD (p=0.02, Hedge’s g=0.82) (**Fig. 3A, Supp Fig. 3A**). The nDNA-encoded CII activity in monocytes also tended to be 51% and 65% higher in MitoD than in controls, consistent with a compensatory upregulation in both MitoD groups. MRC did not differ between MitoD and controls across cell types (**Fig. 3B**).

**Figure 3.**
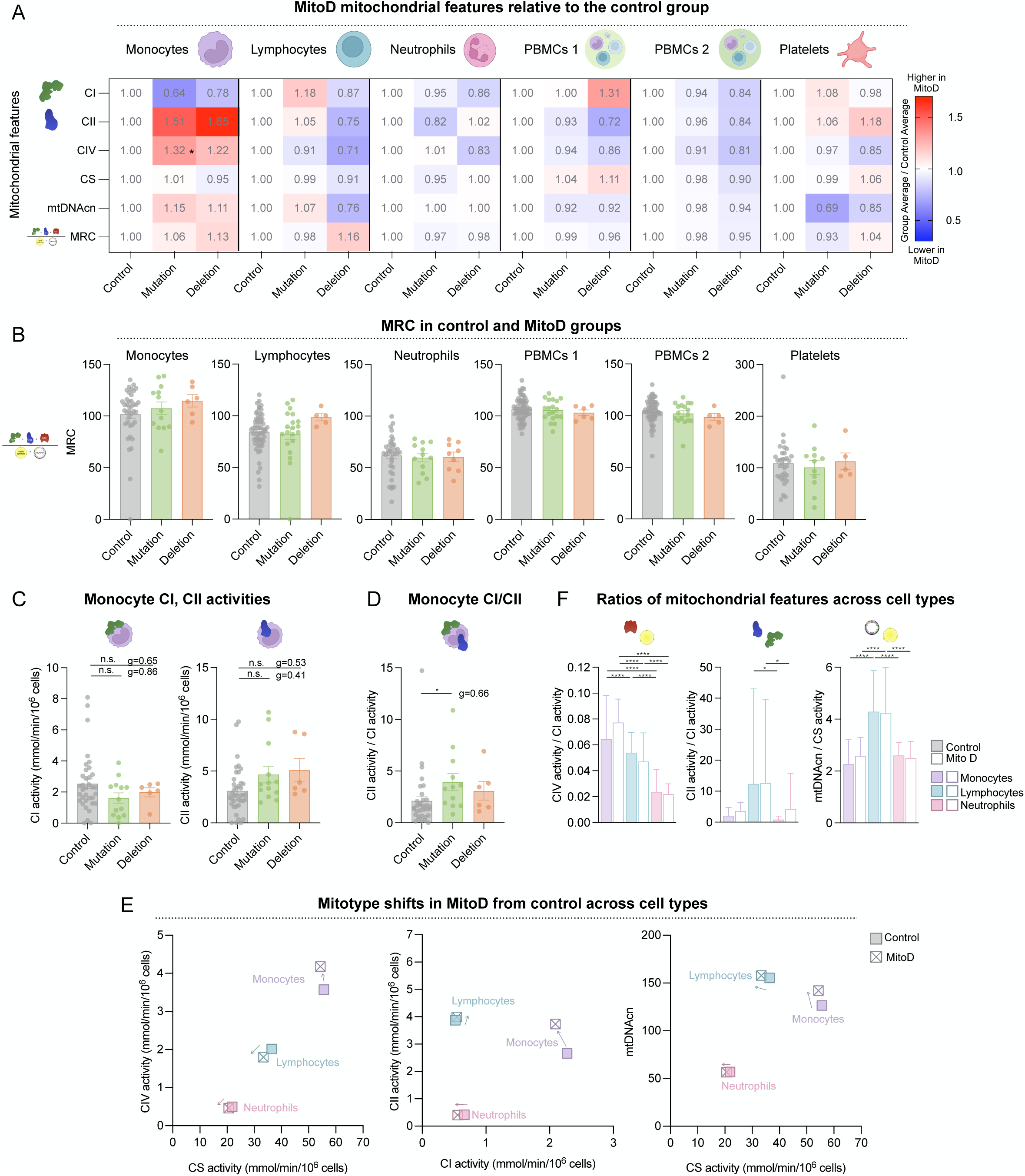
Differences in RC enzyme and CS activity, mtDNA and MRC between control and MitoD across immune cell types, PBMCs and platelets. **(A)** Heatmap of mean mitochondrial measures (CI, CII, CIV, CS activity, mtDNAcn, MRC) in MitoD groups relative to control in immune cell types, PBMCs and platelets. **(B)** MRC activity across different cell types and **(C)** CII activity in monocytes and lymphocytes in healthy control and MitoD groups, individual data points, mean +/- SEM shown. One-way ANOVA, Kruskal Wallis multiple comparison test. *p<0.05, **p<0.01, ***p<0.001, ****p<0.0001. Effect sizes as Hedge’s g (g). **(E)** Bivariate plots and **(F)** ratios showing shift in the relationship between different pairs of mitochondrial enzyme measures and mtDNAcn in immune cell types between healthy controls and MitoD patients. Control n=31-66, Mutation n=11-21, Deletion n=5-13.

Consistent with the biology of mitochondrial disease, mtDNA encoded CI activity in monocytes tended to be 22-36% lower in MitoD patients, and nuclear DNA encoded CII activity was higher by 51-65%. These differences were of medium to large effect sizes (Hedge’s g= 0.41 – 0.86) and the CII/CI activity ratio was 86% higher in MitoD than in controls (p= 0.017, Hedge’s g = 0.66) (**Fig. 3D**). These trends were not found in other cell types (**Fig. 3A**), and neither of the single enzyme measures were significantly different between groups, owing mainly to large inter-individual variance (**Fig. 3C**, see **Supp. Fig 3** for all results).

Overall, the differences in mitochondrial features between control and MitoD were small compared to those between immune cell subtypes (**Fig 3E, Supp. Fig. 3E**), and the significant differences in the ratios seen between cell types of controls were preserved in MitoD (**Fig. 3F**). Differences between cell types were also preserved in MitoD (**Supp. Fig. 3D**), demonstrating cell type specific differences in mitochondrial phenotypes are largely unaffected by mitochondrial diseases.

### Mitochondrial OxPhos capacity does not correlate with mitochondrial disease severity

We next asked whether leukocyte mitochondrial respiratory capacity is related to MitoD biomarkers and symptom severity. **Figure 4A** shows all correlations between immune mitochondrial features and clinical and biomarker metrics, and **Figure 4B-C** illustrates some of these associations using scatterplots. Overall, mitochondrial features were weakly correlated with circulating biomarkers of mitochondrial disease such as growth differentiation factor 15 (GDF15) ^38^ and fibroblast growth factor 21 (FGF21) ^39^. A similar pattern showing only weak and non-significant correlations was observed with clinical measures of symptom severity and functional capacity (r’s= 0.10 – 0.17, **Fig. 4D-F**, see *Methods* for details).

**Figure 4.**
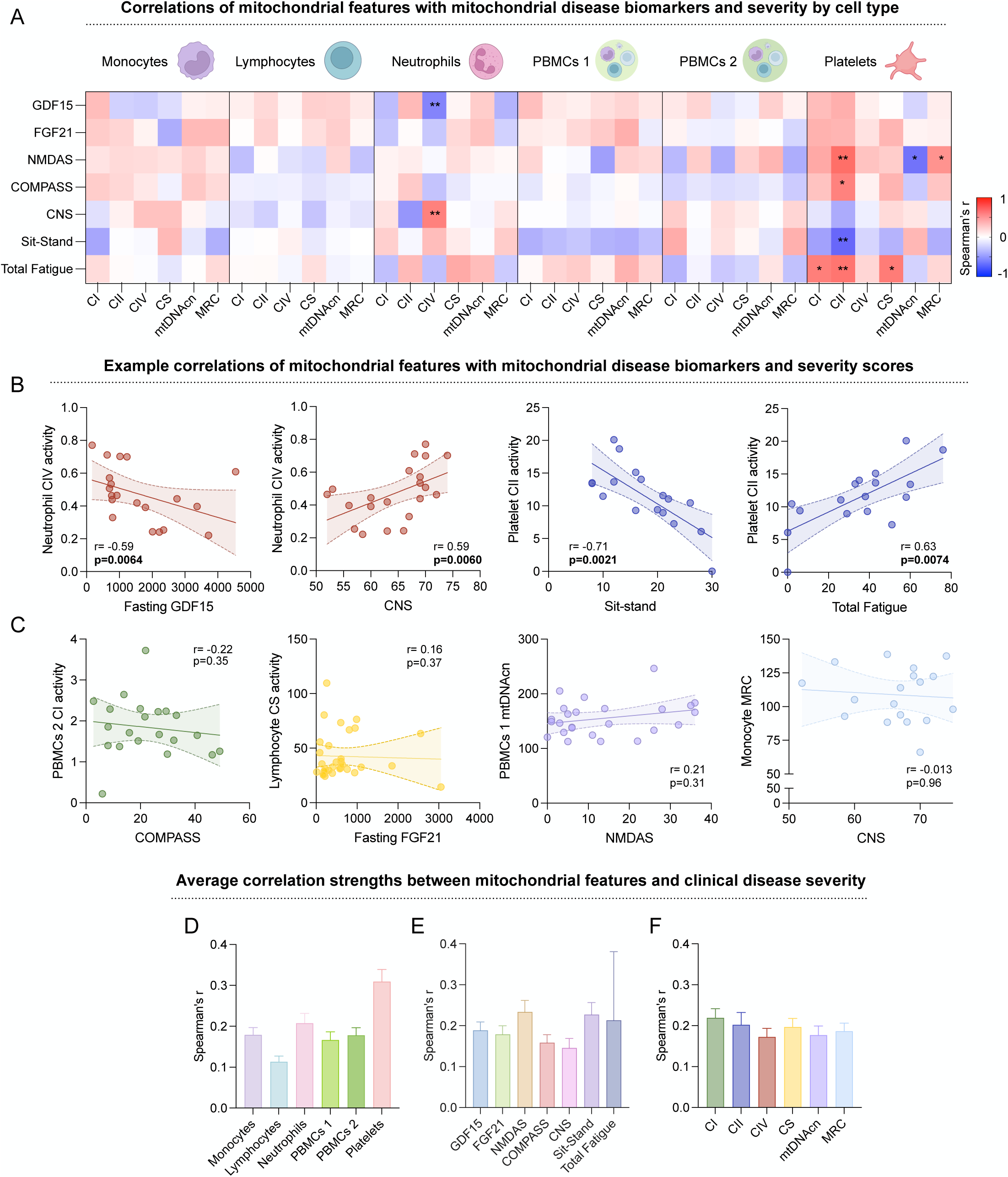
Correlation between clinical biomarkers and mitochondrial disease severity. Correlations between CI, CII, CIV, and CS activity, mtDNA and MRC in immune cell types, PBMCs and platelets and clinical biomarkers and mitochondrial disease severity in mitochondrial disease patients as **(A)** a heatmap, and scatterplots **(B)** showing significant correlations between Neutrophil COX activity and Fasting GDF15, CNS, and between Platelet CII Activity and Sit-stand, total fatigue scores and **(C)** lack of correlation between CI, CII, CIV, and CS activity, mtDNA and MRC in immune cell types, PBMCs and platelets and clinical biomarkers and mitochondrial disease severity in mitochondrial disease patients. Average Spearman correlations (absolute value of r value) between mitochondrial disease biomarkers/severity and mitochondrial features across **(D)** disease biomarker/severity measures by cell type, **(E)** cell types by disease biomarker/severity measure, and **(F)** disease biomarker/severity measures by mitochondrial feature measure. E.g., first bar in (D) represents average correlations across all mitochondrial disease biomarkers/severity and mitochondrial features in monocytes, first bar in (E) represents correlations across all mitochondrial features with GDF15 across all cell types, and first bar in (F) represents correlations across all mitochondrial disease biomarkers/severity with CI activity across all cell types. n=13-33. Spearman’s r labeled on correlation matrix, with significant correlations marked with asterisks in the matrices. *p<0.05, **p<0.01, ***p<0.001, ****p<0.0001. Histograms show mean +/- SEM.

Of all cell types and platelets, platelet mitochondria were the most strongly correlated to clinical measures, and lymphocytes showed the weakest correlations (**Fig. 4D).** Correlations between mitochondrial features and disease severity measures were strongest with NMDAS, but weak overall across disease severity measures (average absolute values of r’s= 0.13 – 0.23) (**Fig. 4E)**. Amongst mitochondrial features, correlations were strongest with CI activity, consistent with MitoD affecting mtDNA encoded complexes; however, also weak overall (average absolute values of r’s = 0.17 – 0.21) (**Fig. 4F)**. This lack of correlation between MRC and disease severity was consistent with the lack of difference in immune cell MRC between MitoD and control groups.

### Changes in mitochondrial biology over time in MitoD and controls

To evaluate the stability of mitochondrial features over different states and time of day, we finally assessed the change in immune mitochondrial biology between the morning/fasted state (Time 1) and post-stress afternoon/fed state (Time 2) (**Fig. 5A**). In healthy controls, CI and CII activities, mtDNAcn and MRC changed significantly from Time 1 to 2 (4% – 25% change, p’s<0.048). Similar changes were observed in MitoD, although only the increase in mtDNAcn from Time 1 to 2 was significant (16% increase, p=0.0039) (**Fig. 5B, D, Supp. Fig 4A, B**). CI activity, however, changed in the opposite direction in MitoD, exhibiting a 13% morning-to-afternoon decline in patients, whereas healthy controls exhibited a significant 17% elevation (p=0.028).

**Figure 5.**
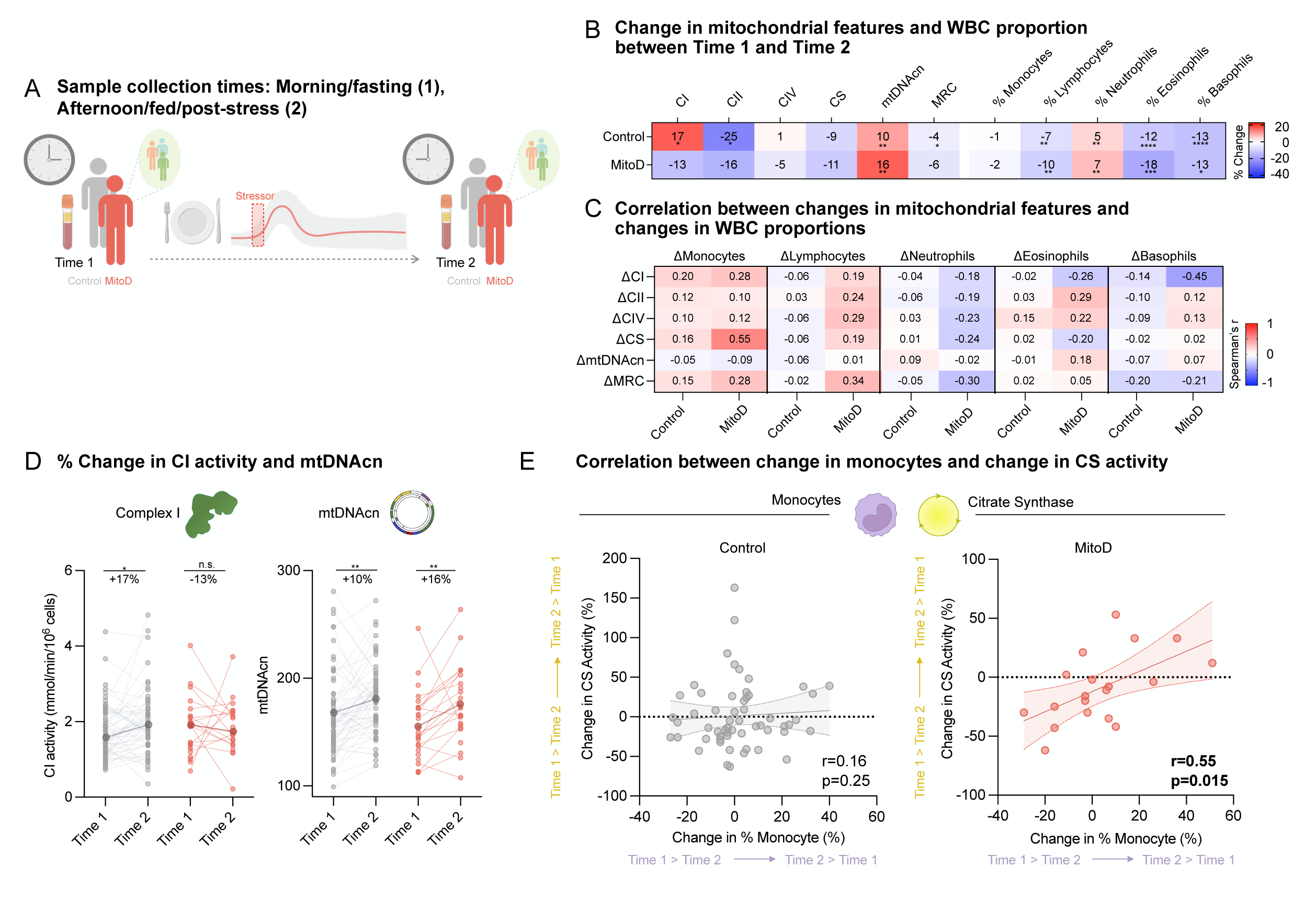
Shift in mitochondrial measures and WBC composition between different timepoints. **(A)** Schematic of sampling time. Time 1 is a morning, fasted sample pre-stress exposure. Time 2 is an afternoon, fed sample, post socio-evaluative stress exposure. Healthy controls are compared to MitoD patients, which include Mutation/MELAS and Deletion groups. **(B)** Median % change in CI, CII, CIV, CS activities, mtDNAcn, MRC in PBMCs, and % composition of WBCs between Time 1 and Time 2 in healthy controls and MitoD patients. **(C)** Correlation between % change in measures of mitochondrial biology and % change in proportion blood cells between Time 1 and Time 2 in healthy controls and MitoD patients. **(D)** Individual and percent changes in CI activity and mtDNAcn in healthy controls (grey) and MitoD patients (red). Raw CI activity and mtDNAcn plotted, median % change (change between Time 1 and 2, relative to Time 1 value) labelled. Darker dot represents average group values at each time point. **(E)** Correlation between change in CS activity and change in % monocytes in Control and MitoD patients. Mitochondrial Features: Control n=58-59, MitoD n=18-19. WBC proportions: Control n=60, MitoD n=32. Spearman’s r labeled on correlation matrices and correlation plots, with significant correlations marked with asterisks in the matrices. *p<0.05, **p<0.01, ***p<0.001, ****p<0.0001. Wilcoxon test for paired timepoint measurements, Mann-Whitney test for group comparisons. *p<0.05, **p<0.01, ***p<0.001, ****p<0.0001.

We also examined if these changes in PBMC mitochondrial features were driven by dynamic changes in cell type composition. The proportion of leukocyte types (lymphocytes, monocytes, neutrophils, eosinophils, and basophils) showed several significant changes from Time 1 to 2 (changes ranged from 5% to 13%, p’s=<0.0061) in all cell types but not in monocytes of healthy controls. MitoD participants displayed similar changes (changes of 2% – 18%, significant p’s= 0.0003 – 0.02, **Fig. 5B, Supp Fig. 4C, D**), suggesting that the gross temporal/diurnal changes in immune biology are not affected in MitoD.

In both control and MitoD, changes in cell type abundance largely did not account for the changes in mitochondrial biology observed (average r = 0.04). One exception was a significant correlation between the change in % monocytes and the change in monocyte CS activity in MitoD (r = 0.55, p = 0.015; Bonferroni corrected p = 0.45), which was not observed in healthy controls (**Fig. 5E**). Further work is required to understand the factors that may explain the temporal changes in mitochondrial OxPhos enzyme activities and mtDNAcn levels and their potential relevance to the immune aspects of mitochondrial diseases.

## DISCUSSION

In this study, we phenotyped OxPhos enzyme activities and markers of mitochondrial content in immune cell subtypes, PBMCs and platelets from dozens of individuals with molecularly defined mtDNA defects causing MitoD. Our results confirm previous work showing distinct mitochondrial profiles of immune cell subtypes ^5, 24, 26, 27^ and demonstrate that these profiles are minimally affected by mtDNA defects in MitoD. Accordingly, the immune cell type-specific mitochondrial phenotypes were preserved in MitoD, did not consistently correlate with MitoD biomarkers or symptom severity, and their temporal variation in mitochondrial phenotypes was only minimally affected, if at all, in MitoD. Together, this work emphasizes the notion that immune mitochondrial biology, although conveniently accessible from minimally invasive blood collection, does not reflect the severity of systemic mitochondrial OxPhos deficiency responsible for clinical signs and symptoms in MitoD.

The distinct cell type-specific mitochondrial phenotypes reported here are similar to previous reports ^24^. Neutrophil mitochondrial features were consistently low across all measures, consistent with previous work demonstrating that their primary role is not ATP synthesis ^27^, and monocytes and PBMCs also displayed consistently higher measures across variables.

Despite the fact that both the m.3243A>G and single large-scale deletions are expected to influence OxPhos complex gene expression and protein synthesis, we note a surprising lack of differences between the mitochondrial OxPhos profiles of participants with MitoD and healthy controls. This result could be explained by the presence of purifying selection in immune cells ^18–21^, where the effect and heteroplasmic levels of these mitochondrial disease-causing mutations are minimized, therefore bringing the OxPhos capacities of MitoD immune cells closer to that of healthy controls.

In alignment with the lack of group differences in leukocyte OxPhos capacity, MRC was not consistently related to mitochondrial disease biomarkers or symptoms. These correlations were weakest in lymphocytes, consistent with lymphoid cells displaying purifying selection, leading to the lowest mutation load across immune cells ^18^. In contrast, platelets displayed the strongest and most significant correlations with mitochondrial disease markers. Purifying selection has yet to be studied extensively in platelets ^18–20^, but given the results here, it could be that megakaryocytes, from which platelets are derived, experience less purifying selection than other immune cell types.

While not significant, monocytes also demonstrated changes in CI and CII activity that are consistent with mitochondrial disease etiology seen in skeletal muscle and brain, with a trend towards lower CI and higher CII average activity ^40, 41^. These monocyte specific changes support the lower level of purifying selection in myeloid cells relative to lymphoid cells ^18^, however further investigation is needed to establish whether these changes are significant and why they are restricted to monocytes. Overall, given that immune cell OxPhos is minimally affected by mitochondrial diseases and not correlated with disease severity, we might expect that immune cell mitochondrial profiles do not provide an accurate representation of somatic (skeletal muscles, brain, etc.) nor whole body OxPhos capacity in MitoD.

In addition to cell-type dependent differences in mitochondrial features, our results reveal potential time-dependent changes in PBMC mitochondrial features. These changes were not explained by changes in immune cell proportions, which was somewhat surprising given the previously established association between the proportion of circulating cell types and PBMC mitochondrial features ^24^. However, additional factors here could have affected mitochondrial phenotypes between these two samples in addition to the time of day – including fasting status and the performance of a stressful socio-evaluative speech task. In previous work, we found that up to 10-15% of variation in mitochondrial OxPhos capacities were associated with subjective reports of positive and negative mood (e.g., feeling energetic and loving) ^42^. Thus, the brief (5-minute) psychosocial stressor that occurs approximately two hours before the afternoon timepoint 2 PBMC sample, and which tends to elicit stress and negative mood, could have contributed to some of the differences observed in OxPhos profiles from the morning/pre-stress to afternoon post-stress timepoint. Whether the causes and consequences of mitochondrial respiratory capacity changes in immune cells reflect that of other organs will also remain an important question to examine. Potential time-dependent changes in mitochondrial biology should also be considered in future studies.

Limitations of this study include the variable sample sizes between cell types. Due to sample collection restrictions, sample size and cell count are limited for certain cell subtypes. This limited the sample size of some correlation analyses, particularly with our limited MitoD population, as well as introducing some noise into our enzyme activity assays. Furthermore, while we demonstrate that immune cell mitochondria are minimally affected in MitoD, the molecular basis for this effect remains unclear. Purifying selection for low heteroplasmy cells in the bone marrow may play a role. Although not possible at this time, assessing the mutation load in each immune cell type alongside with the functional assays would be valuable to establish the extent to which this effect is attributable to mutation load or other non-genetic, humoral factors. But even in the presence of mtDNA heteroplasmy, compensatory mechanisms in specific (sub)populations of immune cells of MitoD patients could, to some extent, mitigate the influence of mtDNA mutations on OxPhos enzyme activities and markers of mitochondrial content.

Overall, this study provides insights into the immune cell-specific and time-dependent PBMC mitochondrial OxPhos profiles of MitoD and healthy controls. Together, the multivariate enzymatic and molecular profiling in multiple leukocyte cell types expands our understanding of immune mitochondrial biology in MitoD. While immune mitochondria are minimally affected by mtDNA mutations and are unlikely to reflect the severity of MitoD or offer reliable diagnostic and/or prognostic information, immune mitochondria could still offer a window into how disease-modifying factors affect mitochondrial biology across organ systems.

## METHODS

### Participant recruitment

Participants were recruited for the Mitochondrial Stress, Brain Imaging, and Epigenetics (MiSBIE) study in adherence to the directives outlined by the New York State Psychiatric Institute IRB protocol #7424, Columbia University Medical Center IRB protocol #AAAU9470, ClinicalTrials.gov NCT04831424^35^. Recruitment occurred from April 2018 to May 2023 at our local clinic at the Columbia University Irving Medical Center and nationally throughout the United States and Canada through clinical partners and Columbia’s RecruitMe site. Recruitment was targeted towards individuals aged 18 to 60, and a total of 110 participants, with 70 healthy controls, (n=48 female, 22 male) and 40 individuals with genetically-defined mitochondrial diseases (MitoD, n=28 female, 12 male) were recruited. In this study, 68 (n=46 female, n=22 male) healthy controls, and 37 (n=25 female, n=12 male) individuals with MitoD were included due to sample collection limitations. All enrolled participants provided written informed consent, authorizing their participation in the investigative procedures and the dissemination of findings.

Healthy controls were eligible if they were willing to provide saliva samples and have a venous catheter installed for blood collection during their hospital visit. Exclusion criteria included cognitive deficits leading to an inability to provide informed consent, recent occurrences of seasonal infections in the four weeks prior to the study, Raynaud’s syndrome, involvement in ongoing trials – therapeutic or exercise-related on ClinicalTrials.gov, the presence of metal inside or outside the body, claustrophobia impeding magnetic resonance imaging (MRI).

MitoD patients were eligible for inclusion if they were genetically diagnosed for either (i) the m.3243A□>□G point mutation, with or without mitochondrial encephalopathy, lactic acidosis, and stroke-like episodes (MELAS), or (ii) a single, large-scale mtDNA deletion-often associated chronic progressive external ophthalmoplegia (CPEO) or Kearns-Sayre syndrome (KSS). Exclusion criteria were similar to that of healthy controls, with additional criteria excluding patients using steroid therapy (oral dexamethasone, prednisone, or similar), or steroid use. Data from 23 patients with the m.3243A>G mutation and 14 patients with single, large-scale mtDNA deletions were included in the present study.

Mitochondrial disease severity quantification and symptom profiling was performed with five approaches: 1) the Newcastle Mitochondrial Disease Scale (NMDAS) ^43^, assessed by a qualified clinician; 2) the Composite Autonomic Symptom Score (COMPASS) ^44^ assessed by a qualified clinician; 3) the Columbia Neurological Score (CNS) minus head circumference assessment, see ^35, 45^ assessed by a qualified clinician; 4) Functional capacity via the 30-second sit-to-stand test (Sit-Stand) ^46^; 5) Perceived fatigue via the Modified Fatigue Impact Scale (Total Fatigue) ^47^.

A socio-evaluative speech task was conducted on all participants as described previously ^35^. Briefly, study participants were asked to prepare and deliver a speech to a court judge defending themselves against alleged transgression, shoplifting. Participants had two minutes to prepare their speech and three minutes to deliver their speech in front of an intimidating evaluator wearing a white coat. Participants were told their performance was being filmed for later evaluation and during the stress task they could see themselves in a full-length mirror. An intravenous catheter was placed in the right arm at least 45 minutes before the beginning of the speech task to minimize the effect of any discomfort experienced by the participants related to blood collection on the measured parameters.

### Sample Collection

Blood was collected via a central venous catheter both at ∼10 am while participants were in a fasted state, and in the afternoon, around 4 pm 120 minutes after the onset of the socio-evaluative speech task. Five 8.5mL ACD-A tubes (BD-364606) were collected at ∼10 am from the left arm for the morning PBMC samples, platelets and specific immune cell subtypes. Immediately after blood collection, samples are inverted 10-12 times, and placed in a Styrofoam box at room temperature for transport to the laboratory (5-minute walk). Tubes were then centrifuged at 500g for 15 minutes at room temperature with the brakes off. For the afternoon PBMC sample, blood was collected via an intravenous catheter in the antecubital vein of the left arm at ∼4 pm and collected in one 8.5ml tube ACD-A(BD-364606). The ACD-A tube was used for PBMC isolation, and was transported to the laboratory at room temperature, where it was centrifuged at 500g for 15min at room temperature with the brakes off.

#### Platelet Isolation

Morning fasting tubes were processed for platelet isolation. 3ml of plasma were isolated from each ACD-A tube and pooled in a 15ml conical centrifuge tube with 15μl of 1uM prostaglandin I2 (PGI2) (Cayman Chemicals, #18220) to prevent platelet aggregation. Red blood cells were pelleted with a 1000g spin for 5min at room temperature, and then the supernatant was transferred to a new 15ml tube and centrifuged for 1500g for 10min at room temperature. The supernatant was discarded again, and the pellet resuspended, before 10ml of PBS-PGI2 (1mM) was added and mixed into the cell suspension until homogenous. The total volume was centrifuged at 1500g for another 10min, supernatant was discarded, and the remaining platelets were resuspended in 3ml PBS-PGI2 (1M) buffer. Platelets were quantified with a turbidity assay for the first 28 participants, estimated using the formula described in ^48^:

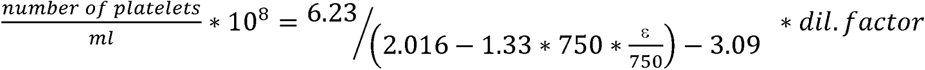

and the remaining samples were quantified using a Bicinchoninic acid (BCA) assay (Bioworld #20831001) for whole protein content. Samples were stored in liquid nitrogen until sample homogenization and enzyme activity assays were run.

#### PBMC isolation

Peripheral blood mononuclear cells (PBMCs) were isolated from morning fasting and afternoon post-stress blood. For the morning sample, after collection of plasma from each ACD- A tube for platelet isolation, the first ACD-A tube was collected in a 15ml conical tube and used to isolate PBMCs. For the afternoon sample, one ACD-A tube was set aside for PBMC isolation. After collection of plasma, blood from the ACD-A tube was transferred to fresh 15ml conical tube. 5ml HBSS was added to the ACD-A tube to wash and collect any remaining cells and transferred to the 15ml conical tube. The diluted blood was aspirated and slowly overlayed over 4mL of preloaded Histopaque 1077 (Sigma, # 10771) in a 15ml conical tube. The tube was then spun at 400g for 30min at room temperature with brakes off. The cell layer was transferred to a 15ml conical tube with 2ml HBSS, and then HBSS was added until the final volume reached 15ml. The tube was centrifuged at 500g for 10min to pellet cells, supernatant was discarded, and cells resuspended with HBSS to 15ml for washing. The tube then underwent 2 cycles of centrifugation at 200g for 10min at room temperature, supernatant discard and resuspension with 15ml HBSS. After the third wash, the supernatant was discarded, and cells were resuspended in 1ml of HBSS. 10μl of suspension were mixed with 10μl of trypan blue for counting in the Countess II (ThermoFisher Scientific, AMQAF1000). Total cell count in the suspension was estimated via the cell count and recorded, samples were aliquoted before spinning at 2000g for 2min, having the supernatant aspirated, and stored in a −80C freezer before transfer into liquid nitrogen.

#### Immune cell subtype isolation

To purify immune cell subtypes (monocytes, lymphocytes and neutrophils), 3ml of buffy coat was aspirated from the remaining four ACD-A tubes after platelet and PBMC isolation and pooled in a 50ml conical tube following plasma aspiration. The tube was filled to 50ml with HBSS and inverted to mix. 25ml of this leukocyte cell suspension was added to each of two 50ml Histopaque double gradient tubes (10ml Histopaque 1077 and 10ml Histopaque 1119), and spun at 700g for 30min at room temperature, with the brakes off. The top diluted plasma layer was removed, and the middle mononuclear (MCN) cell layer was collected and pooled in a 50ml tube with 5ml HBSS. The polymorphonuclear (PMN) cell layer was collected and pooled in a 50ml tube with 5ml HBSS. These tubes were centrifuged at 700g for 10min at room temperature, and supernatant was discarded. Cell pellets were resuspended, and 1ml of HBSS/BSA (0.5% BSA in HBSS) (Sigma, #A3733) was added to each tube. Cells were collected and the cell suspension mixed, and aliquoted into two 1.5ml tubes each. These tubes then were centrifuged at 700g for 40 seconds to pellet cells, supernatant was discarded, and pellets were resuspended. 250μl of HBSS/BSA was added to dissolve cell aggregates.

Monocyte, neutrophil and lymphocytes were isolated using magnetic bead tagged antibodies and MACs separator columns (Miltenyi Biotech, #130042401). Monocytes were isolated using 66.4μl of magnetic bead labelled CD14 antibody (Miltenyi Biotec, #130050201) from the MCN tube and the neutrophils were isolated using 66.4μl of magnetic bead labelled CD15 antibody (Miltenyi Biotec, #130046601) from the PMN tube. Lymphocytes were isolated from the flow through after the monocyte isolation. The flowthrough was centrifuged at 300g for 10min, the supernatant was discarded, the cell pellet was resuspended in 1ml of HBSS/BSA, and the suspension was collected. The tube was then centrifuged at 700g for 40 seconds, and negative selection of lymphocytes was performed using 66.4μl each of magnetic beads labeled CD61(Miltenyi Biotec, #130051101) and CD235 (Miltenyi Biotec, #130050501) antibodies. Cells were isolated after incubating the cell suspension with antibody for 15min on wet ice, equilibrating MACs separator LS columns with 3ml HBSS/BSA, adding 1ml HBSS/BSA to the cell suspension, and centrifuging the tube at 700g for 40 seconds. The supernatant was discarded, and cell resuspended with 1ml HBSS/BSA. The cell suspension was added to the LS column and washed with 3ml HBSS/BSA three times, before collection of cells with 5ml of HBSS. The collected cells were centrifuged at 100g for 10min at room temperature to pellet cells. The supernatant was discarded, and cell pellets resuspended. Cell counts were performed with the Countess II, and stored in liquid nitrogen until enzymatic assays were run.

In this study, a total of 435 samples were assayed across all cell types: 92 morning fasting PBMC (PBMCs 1), 84 afternoon post-stress PBMC (PBMCs 2), 59 monocyte, 94 lymphocyte, 57 neutrophil, and 49 platelet samples. Counts per cell type per group are included in the below table. Samples of each cell type were run on the same plate, except for one morning fasting PBMC and one neutrophil sample that were run on the same plate as the platelets. Each plate was run with 1-3 mouse reference samples, in duplicate, for batch correction.

| Cell type | Control | m.3243A>G | Deletion | Total |
| --- | --- | --- | --- | --- |
| PBMCs 1 | 55 | 23 | 14 | 92 |
| PBMCs 2 | 51 | 20 | 13 | 84 |
| Monocytes | 39 | 12 | 8 | 59 |
| Lymphocytes | 58 | 22 | 14 | 94 |
| Neutrophils | 33 | 15 | 9 | 57 |
| Platelets | 28 | 13 | 8 | 49 |

### Mitochondrial Enzymatic Assays

#### Homogenization

Cell pellets were homogenized for enzymatic activities as previously described ^24^. Homogenization was completed in 500μl of homogenization buffer 1mM EDTA, 50mM Triethanolamine) with two tungsten beads (Qiagen Cat#69997) in a Tissue Lyser (Qiagen, Cat# 85300) with pre-chilled racks. Two rounds of 30 cycles/s for 1min were completed, with incubation of samples at 4°C in ice for 5min in between rounds. Samples were vortexed before use to ensure their homogeneity. Mouse reference samples were prepared by homogenizing mouse tissue at a ratio of 1mg tissue to 180μl homogenization buffer using the same procedure as the cells, but with an additional 3 freeze/thaw cycles. Mouse tissue homogenates were pooled to produce three unique reference samples.

#### Enzyme activity assays

Colorimetric enzyme activity assays for complex I (NADH-ubiquinone oxidoreductase, CI), complex II (succinate-ubiquinone oxidoreductase, SDH), complex IV (cytochrome c oxidase, COX) and citrate synthase (CS) were measured spectrophotometrically as previously described (Rausser et al., 2021), with minor modifications. Enzymatic assays were performed in 96-well plates and recorded on a Spectramax M2 (Spectramax Pro 6, Molecular Devices), using the following volumes of homogenate: Complex I: 15μl, COX and SDH: 20μl, and CS: 10μl.

##### Complex I (CI)

Total CI activity was determined by measuring the change in absorbance at 600 nm over 10min, in a 100uM potassium phosphate reaction buffer (pH 7.5) containing 2mM EDTA, 3.mg/ml BSA, 0.25mM potassium cyanide, 10μM decylubuquinone, 100μM DCIP, 200μM NADH, and 0.4μM antimycin A. Non-specific activity was detected in the presence of 500uM rotenone and 200uM piericidin A, specific inhibitors of CI.

##### Complex II (CII)

Total CII activity was determined by measuring the change in absorbance at 600nm over 15min, in a 100mM potassium phosphate reaction buffer (pH 7.5) containing mM EDTA, 1mg/ml BSA, 4μM rotenone, 1mM succinate, 0.25mM potassium cyanide, 100μM decylubuquinone, 100μM DCIP, 200μM ATP, and 0.4μM antimycin A. Non-specific activity was measured in the presence of malonate (5 mM), a specific inhibitor of CII.

##### Complex IV (CIV)

Total CIV activity was determined by measuring the change in absorbance at 550nm over 20min for lymphocytes, neutrophils, monocytes and platelets, and 10min for PBMCs, in a 100mM potassium phosphate reaction buffer (pH 7.5) containing 0.1% n-dodecylmaltoside and 50μM of purified reduced cytochrome *c*. Spontaneous cytochrome *c* oxidation was determined by measuring change in absorbance without cell lysate.

##### Citrate Synthase (CS)

Total CS activity was determined by measuring the change in absorbance at 412nm over 8min, in a reaction buffer (200mM Tris, pH 7.4) containing 0.2mM acetyl-CoA 0, 0.2mM 5,5’-dithiobis-(2-nitrobenzoic acid) (DTNB), 0.55mM oxaloacetic acid, and 0.1% Triton X-100. Non-specific activity was measured in the absence of oxaloacetate from the reaction buffer.

Total and non-specific activities for each enzyme were assessed in triplicate at 30°C. The number of replicates of spontaneous cytochrome *c* oxidation measure was equal to the number of samples assessed on for total CIV activity. Activities were determined using the change in absorbance over time for each enzyme, where the trace for absorbance vs. time was linear and maximal. To transform change in OD into enzymatic activity, the following molar extinction coefficients were used: DTNB – 13.6Lmol^−1^cm^−1^, reduced cytochrome c – 29.5Lmol^−1^cm^−1^, DCIP – 16.3Lmol^−1^cm^−1^.

In the present study, PBMC activity values were lower relative to other cell types than previously reported ^24^, likely as a result of a more thorough platelet depletion, resulting in lower platelet contamination in PBMCs.

#### Specific activity calculation, replicate cleaning

Specific activities were calculated by taking the average of total activities and subtracting the average non-specific/spontaneous activities. Any specific activity that was calculated to be <0 was deemed undetectable and replaced by 0. This was done for 3% of all enzyme activity measurements. For samples that had a CV greater than the following cut-off CVs within the replicates of measure and did not have data excluded due to non-linear traces, the measure/s that were more than 1 standard deviation from the mean of the replicates were excluded. If any data was excluded due to non-linear traces, the average of the remaining measures was taken. The cut off CVs are as follows: CS, CI, SDH – 15%, COX – 35% for PBMCs and 30% for all other cell types.

#### Batch correction

Batch correction for each enzyme assay was performed by dividing each enzyme activity per plate by a correction factor. The correction factor was calculated by taking average activity of each mouse reference sample on each plate normalized to the average of the reference sample across all plates. This was value averaged across the reference samples for a given plate to produce a correction factor. The correction factors are as follows:

| Enzyme | PBMCs 1 | PBMCs 2 | Monocytes | Lymphocytes | Neutrophils | Platelets |
| --- | --- | --- | --- | --- | --- | --- |
| CS | 1.01 | 0.98 | 0.98 | 0.97 | 0.96 | 1.08 |
| CII | 1.03 | 0.94 | 1.00 | 0.85 | 0.77 | 0.87 |
| CII | 0.54 | 0.85 | 1.44 | 0.64 | 0.93 | 1.2 |
| CIV | 1.00 | 1.04 | 0.95 | 0.78 | 0.91 | 1.15 |

### qPCR

#### qPCR assay

mtDNA copy number (mtDNAcn) and nuclear DNA (nDNA) were measured via qPCR as previously described (Rausser et al., 2021), with minor adjustments. 20μl of the same homogenate used for enzymatic activity measurements was lysed in 180μl lysis buffer (6% Tween20 (Sigma #P1379), 114 mM Tris-HCl pH 8.5 (Sigma #T3253), and 200 µg/mL Proteinase K (Thermofisher #AM2548) for 16h at 55°C, followed by Proteinase K inactivation at 95°C for 10min, and were maintained at 4°C until used for qPCR if qPCR was done within 24h of lysis, or stored at −80°C until qPCR was performed if the assay was performed more than 24h from lysis. For each primer pair, qPCR reactions were performed in two plates of triplicates in 384-well qPCR plates with 8μl of lysate and 12μl of master mix, cycled in a QuantStudio 7 flex qPCR instrument (Applied Biosystems, Cat# 4485701) with the following cycling conditions: 50°C for 2min, 95°C for 20s, followed by 40 cycles of 95°C for 1 s, and 60°C for 20min (total run time: 40min).

Taqman chemistry was used to quantify mitochondrial and nuclear amplicons simultaneously for two pairs of primers, ND1/B2M and COX1/RNaseP. Master Mix1 for ND1 (mtDNA) and B2M (nDNA) included: TaqMan Universal Master mix fast (Life Technologies #4444964), 300nM of primers and 100nM probe (ND1-Fwd: GAGCGATGGTGAGAGCTAAGGT, ND1-Rev: CCCTAAAACCCGCCACATCT, Probe: HEX-CCATCACCCTCTACATCACCGCCC-3IABkFQ. B2M-Fwd: CCAGCAGAGAATGGAA AGTCAA, B2M-Rev: TCTCTCTCCATTCTTCAGTAAGTCAACT, Probe: FAM-ATGTGTCTGGGT TTCATCCATCCGACA-3IABkFQ). Master Mix2 for COX1 (mtDNA) and RNaseP (nDNA) included: TaqMan Universal Master Mix fast, 300nM of primers and 100nM probe (COX1-Fwd: CTAG CAGGTGTCTCCTCTATCT, COX1-Rev: GAGAAGTAGGACTGCTGTGATTAG, Probe: HEX-TGCC ATAACCCAATACCAAACGCC-3IABkFQ. RNaseP-Fwd: AGATTTGGACCTGCGAGCG, RNaseP-Rev: GAGCGGCTGTCTCCACAAGT, Probe: FAM-TTCTGACCTGAAGGCTCTGCGCG-3IABkFQ).

#### mtDNAcn calculation

mtDNAcn was calculated using the ΔCt between the nDNA and mtDNA Cts. The ΔCt was calculated for both the ND1/B2M and COX1/RNaseP primer pairs by subtracting the average nDNA Ct by the average mtDNA Ct for each example. mtDNAcn was calculated for each primer pair by raising 2 to the ΔCt multiplied by two to account for the diploid nuclear genome (*mtDNAcn* = 2*ΔCt* * 2).

#### mtDNAcn data cleaning

The mean and median deviation between the mtDNAcn calculated using ND1/B2M and COX1/RNaseP primer pairs were 50.3% and 51.6%, respectively (higher in ND1/B2M), as expected. The average of both primer/probe sets is taken as the closest estimate of mtDNAcn for each sample. Samples whose mtDNAcn deviation was more than 20% away from the mean deviation of the plate were systematically checked for clear amplification failures in any primers. If there was a clear amplification failure, the mtDNAcn calculated using the failed primer was replaced with a predicted value based on a plate-specific linear regression model. In these samples, COX1 was the only failed primer. Those without a clear amplification failure had their two mtDNAcn values averaged as per the other samples.

#### mtDNA in platelets

Platelets do not have nDNA, we report mtDNA content normalized to protein concentration derived from the linearized mtDNA Ct divided by protein concentration of each sample.

### BCA protein assay

A Bicinchoninic acid (BCA) assay was run to measure total protein content for normalization of the enzymatic activities to protein concentration in platelets. A BCA assay kit (Bioworld #20831001) was used, with 1:50 Reagent A: B. Bovine serum albumin (BSA) protein standards were prepared using a stock 50mg/ml BSA (Sigma A3733) solution. Standards were diluted in homogenization buffer via serial dilution to produce a standard curve with the following concentrations: 1000ug/ml, 500ug/ml, 250ug/ml, 125ug/ml, 62.5 ug/ml, 31.25ug/ml, 15.62ug/ml.

The assay was run with a 1/2 dilution of the sample homogenate in homogenization buffer. The samples were run in duplicate, with each duplicate on one of two plates, with 25μl of diluted sample or standard added to each well. The assay was run with 2 standard curves, which were averaged for interpolation of protein concentration. Samples with CVs between duplicates greater than 20% were rerun. After reruns, if CVs between replicates were not significantly improved, the average of all 4 values was taken. If one of the 4 values was a clear outlier and the other 3 agree, the outlier was removed. Any values with a negative interpolated protein concentration were removed.

### Normalization of Enzyme activity

To account for differences in cell number and protein concentrations in each sample, the following normalization methods were performed.

#### nDNA normalization

Normalization of enzyme activities to nDNA was performed by dividing the enzyme activities by an nDNA adjustment factor. This nDNA adjustment factor was calculated by dividing the linearized nDNA Ct of the sample by the linearized nDNA factor for all samples, excluding platelets. The linearized nDNA factor was calculated as 2^nDNA^ ^Ct,^ where nDNA Ct is the average of the B2M and RNaseP Cts. nDNA normalization was done for PBMCs, monocytes, neutrophils, and lymphocytes.

#### Protein concentration normalization (platelets)

Normalization of enzyme activities to protein concentration for platelets was performed by dividing the enzyme activities by a protein adjustment factor. This protein adjustment factor was calculated by dividing protein concentration of the individual sample by the average of protein concentrations for all platelets.

### Outlier exclusion

Outliers that were not biologically plausible were systematically identified per cell type and enzyme combination after normalization to nDNA (all excluding platelets) and protein concentration (platelets only). Samples that were more than 3 times the inter-quartile range (IQR) higher than the 3^rd^ quartile, and 3 times the IQR lower than the 1^st^ quartile were identified as outliers and removed from the cleaned dataset.

All data processing from the raw absorbance vs. time data to the final data upon which analyses were conducted were performed in R. Code for analyses is available at https://github.com/mitopsychobio/2024_Immune_MRC_Liu.

### Mitochondrial Respiratory Capacity (MRC)

The Mitochondrial Respiratory Capacity (MRC) considers the dimensional scaling of each variable. RC enzyme activities should scale with the two-dimensional surface area of the mitochondrial inner membrane, and mitochondrial content measures should scale with the three-dimensional volume of the mitochondrial matrix. MRC is calculated from the average of the square root of RC enzyme activities divided by the average of the cube root of mitochondrial content measures: 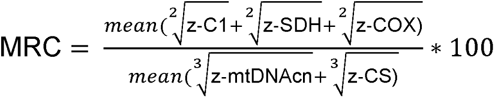. Each each variable was mean-centered (divided by the mean) to to harmonize the effects of each component on the final metric. MRC was calculated for each participant using the nDNA adjusted enzyme values for all cell types excluding platelets, and using the protein adjusted enzyme values for the platelets.

The mean centered values used to calculate MRC for PBMCs, lymphocytes, monocytes and neutrophils were calculated using the averaged values across all these cell types, and the mean centered values used to calculate MRC for platelets were calculated using the averaged platelet values.

### Complete blood count (CBC) with differential

Complete blood count (CBC) with differential was evaluated from blood collected in the fasting state at 10am and in the afternoon 120min after the onset of the socio-evaluative speech task and used to determine immune cell percentages between the two timepoints in this study. CBC with differential was performed with a Sysmex XN in an FDA-approved, and CLIA-certified laboratory. Additional information about the assay can be found at https://www.testmenu.com/nyphcolumbia/Tests/624139

### Effect of time

In previous studies, samples collected more recently (i.e., stored for shorter duration before the mitochondrial assays) had higher activities than those collected earlier in the study when samples were stored at −80°C. This is likely due to the time-dependent degradation of enzyme activities during the storage period. In this MiSBIE dataset however, with liquid nitrogen sample storage, we did not observe a consistent significant decrease in enzyme activity over time across cell types.

The association between time and enzymatic activities are as follows (slope labelled is change in activity over time in days of the linear regression model). The data shown is the nDNA-adjusted data for all cell types except platelets, and protein concentration adjusted data for platelets. (Supp. Fig. 5)

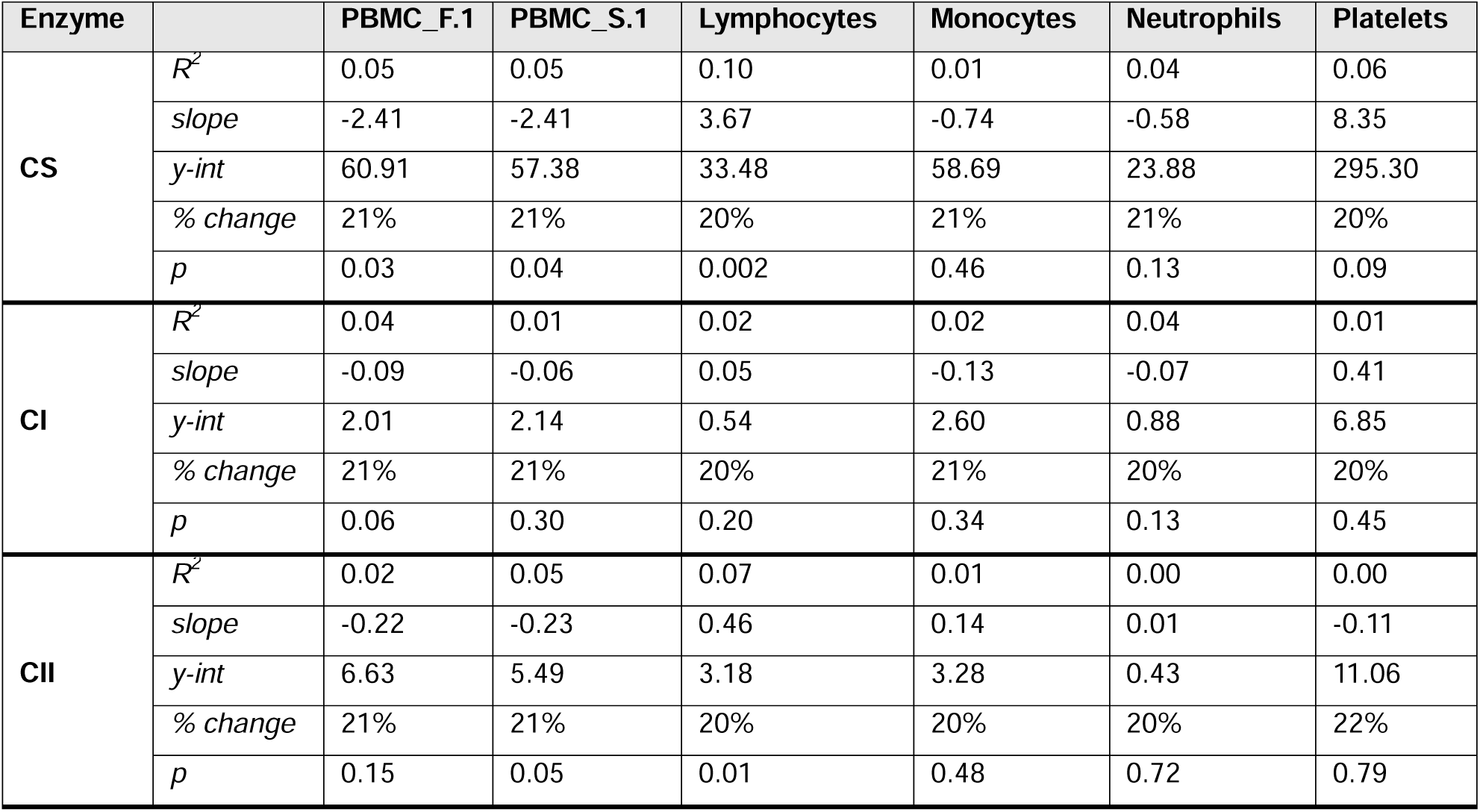
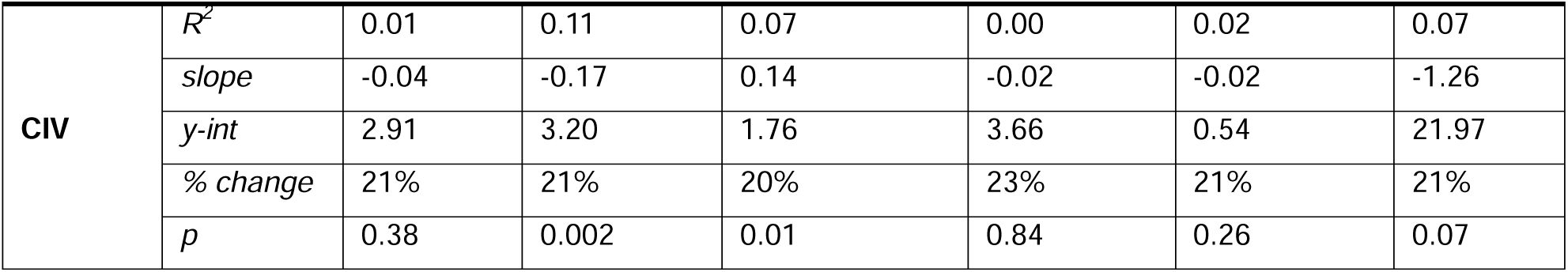

| Enzyme |  | PBMC_F.1 | PBMC_S.1 | Lymphocytes | Monocytes | Neutrophils | Platelets |
| --- | --- | --- | --- | --- | --- | --- | --- |
| CS | $R^2$ | 0.05 | 0.05 | 0.10 | 0.01 | 0.04 | 0.06 |
|  | slope | -2.41 | -2.41 | 3.67 | -0.74 | -0.58 | 8.35 |
|  | y-int | 60.91 | 57.38 | 33.48 | 58.69 | 23.88 | 295.30 |
|  | % change | 21% | 21% | 20% | 21% | 21% | 20% |
|  | p | 0.03 | 0.04 | 0.002 | 0.46 | 0.13 | 0.09 |
| CI | $R^2$ | 0.04 | 0.01 | 0.02 | 0.02 | 0.04 | 0.01 |
|  | slope | -0.09 | -0.06 | 0.05 | -0.13 | -0.07 | 0.41 |
|  | y-int | 2.01 | 2.14 | 0.54 | 2.60 | 0.88 | 6.85 |
|  | % change | 21% | 21% | 20% | 21% | 20% | 20% |
|  | p | 0.06 | 0.30 | 0.20 | 0.34 | 0.13 | 0.45 |
| CII | $R^2$ | 0.02 | 0.05 | 0.07 | 0.01 | 0.00 | 0.00 |
|  | slope | -0.22 | -0.23 | 0.46 | 0.14 | 0.01 | -0.11 |
|  | y-int | 6.63 | 5.49 | 3.18 | 3.28 | 0.43 | 11.06 |
|  | % change | 21% | 21% | 20% | 20% | 20% | 22% |
|  | p | 0.15 | 0.05 | 0.01 | 0.48 | 0.72 | 0.79 |
| **CIV** | $R^2$ | 0.01 | 0.11 | 0.07 | 0.00 | 0.02 | 0.07 |
|  | *slope* | -0.04 | -0.17 | 0.14 | -0.02 | -0.02 | -1.26 |
|  | *y-int* | 2.91 | 3.20 | 1.76 | 3.66 | 0.54 | 21.97 |
|  | *% change* | 21% | 21% | 20% | 23% | 21% | 21% |
|  | *p* | 0.38 | 0.002 | 0.01 | 0.84 | 0.26 | 0.07 |

### Statistical Analyses

All statistical analyses were conducted using GraphPad Prism (Version 10.2.0). Effect sizes were computed as Hedge’s g (g). Mann-Whitney t-tests were used to compare differences in mitochondrial features at different timepoints. Spearman’s rank (r) correlations were used to compute associations between cell types, between mitochondrial features, between mitochondrial features and disease severity measures, and between mitochondrial features and WBC proportions. One-way non-parametric ANOVA Kruskal-Wallis tests were used to compare mitochondrial features between cell types and disease groups.

## Data Availability Statement

Anonymized data not published in the article will be shared upon reasonable request.

## Author contributions

M.P., M.H., C.T., and R.P.J. designed the MiSBIE study. C.K. recruited and enrolled MiSBIE participants. M.K. and A.S.M processed samples. M.K. assisted with enzyme activity assays. M.H. performed the medical examinations. A.S.M and C.T. advised on data processing. C.C.L. performed enzyme activity assays, qPCR assays, data processing and data analysis. C.C.L and M.P. drafted the manuscript. M.P. obtained funding and supervised the project. All authors reviewed the final version of the manuscript.

## Acknowledgements

Work of the authors is supported by R01MH122706, R01AG066828, R01MH119336, the Wharton Fund, and the Baszucki Brain Research Fund to M.P.

## SUPPLEMENTAL FIGURE LEGENDS

**Figure S1.**
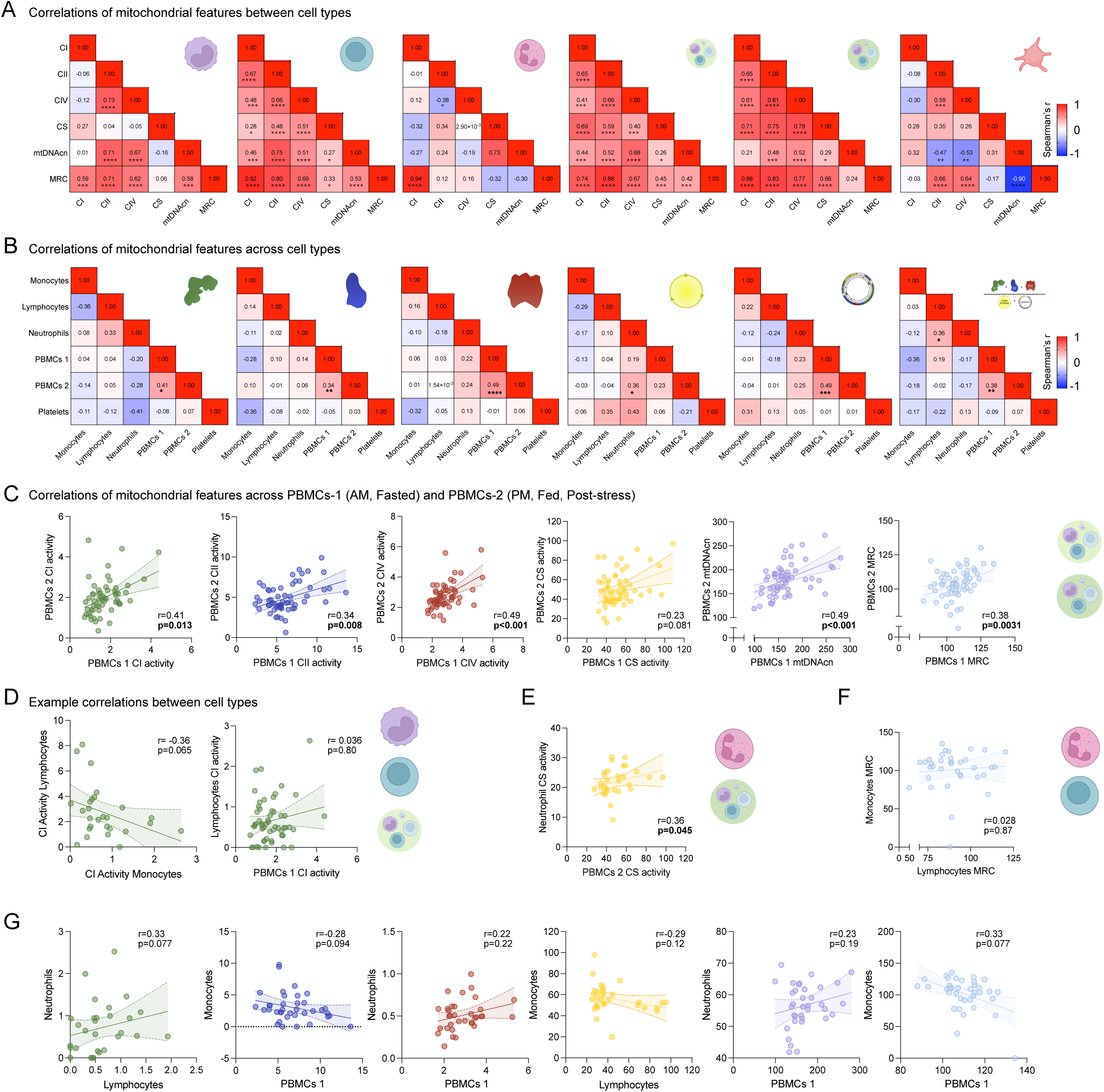
Correlations between mitochondrial measures within and between cell subtypes for mitochondrial enzyme activities, mtDNAcn, and MRC in healthy controls. Correlations **(A)** between mitochondrial enzyme activities, mtDNAcn, and MRC in Monocytes, Lymphocytes, Neutrophils, PBMCs and platelets across cell types and **(B)** of CI, CII, CIV, and CS activity, mtDNA and MRC between immune cell types, PBMCs and platelets. **(C)** Correlation between CI, CII, CIV, and CS activity, mtDNA and MRC of PBMCs collected at different timepoints for each participant and their respective linear regression line with 95% confidence interval of the best fit line. Enzyme activity measurements in mmol/min/10^6^ cells. Correlations between **(D)** CI activity of different cell types showing lack of significant correlations, **(E)** CS activity of Time 2 PBMCs and Neutrophils, (F) MRC of Lymphocytes and Neutrophils, **(D)** CI, CII, CIV, and CS activity, mtDNAcn and MRC in different cell types showing lack of significant correlations. n=31-66. Non-parametric Spearman’s correlations. Spearman’s r labeled on correlation matrices and correlation plots, with significant correlations marked with asterisks in the matrices. *p<0.05, **p<0.01, ***p<0.001, ****p<0.0001.

**Figure S2.**
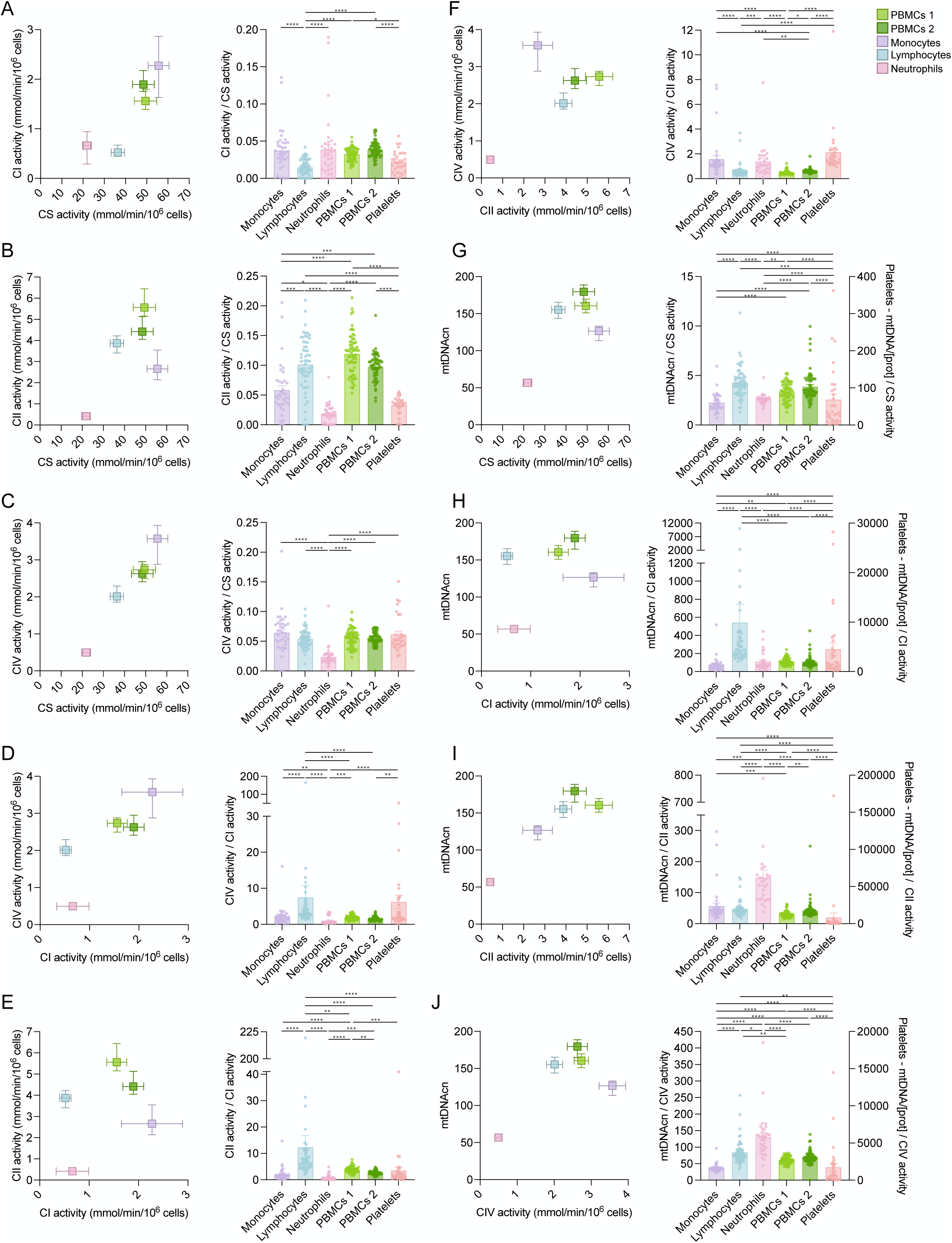
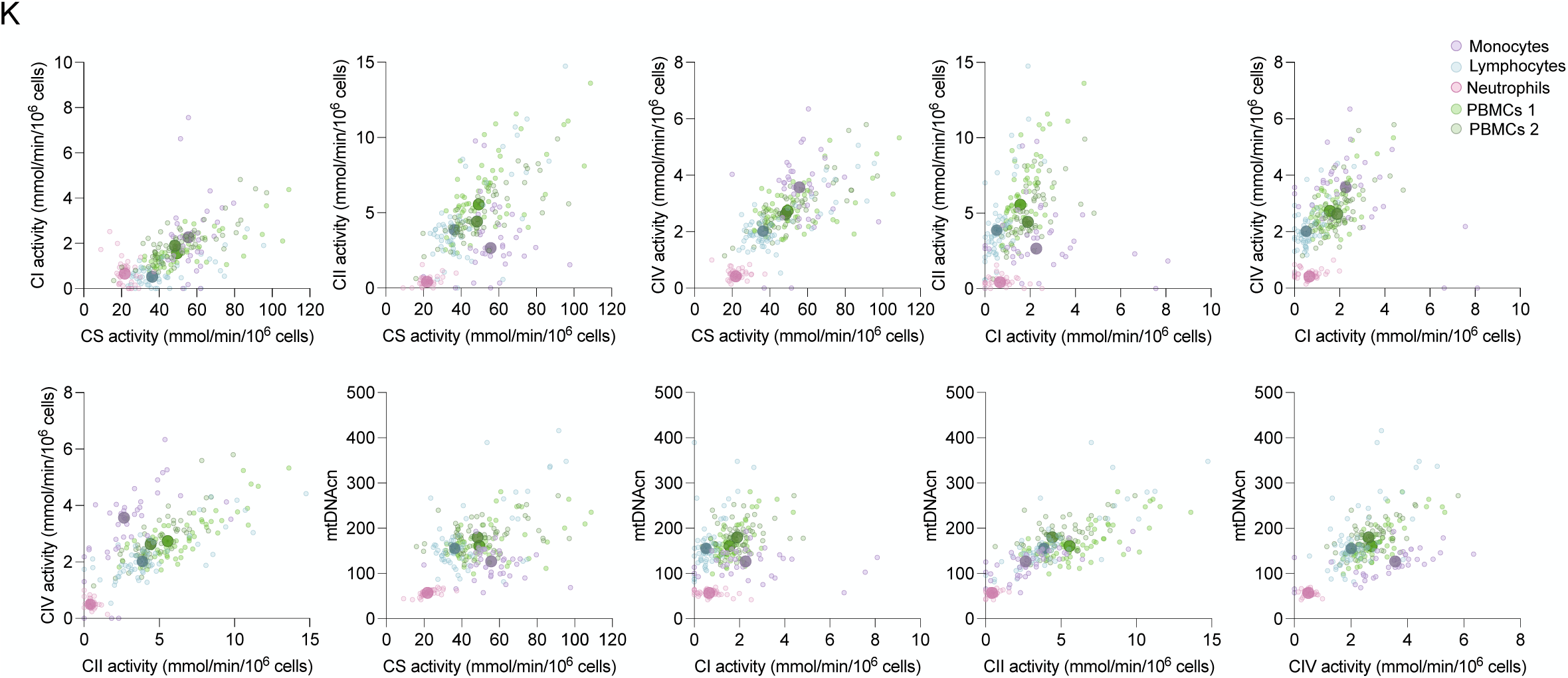
Relationships between different measures of mitochondrial biology by cell type. **(A)** CS vs. CI, **(B)** CS vs. CII, **(C)** CS vs. CIV, **(D)** CI vs. CII, **(E)** CI vs. CIV, **(F)** CII vs. CIV, **(G)** CS vs. mtDNAcn, **(H)** CI vs. mtDNAcn, **(I)** CII vs. mtDNAcn, and **(J)** CIV vs. mtDNAcn and ratio in different cell types. Bivariate plots show median +/- 95% confidence interval. Histograms show individual values, with mean +/- SEM. One-way ANOVA, Kruskal Wallis multiple comparison test. *p<0.05, **p<0.01, ***p<0.001, ****p<0.0001. Individual data points, mean +/- SEM shown. **(K)** Bivariate plots showing relationship between different pairs of mitochondrial enzyme measures and mtDNAcn in different cell types of healthy controls for individual participants. n=31-66.

**Figure S3.**
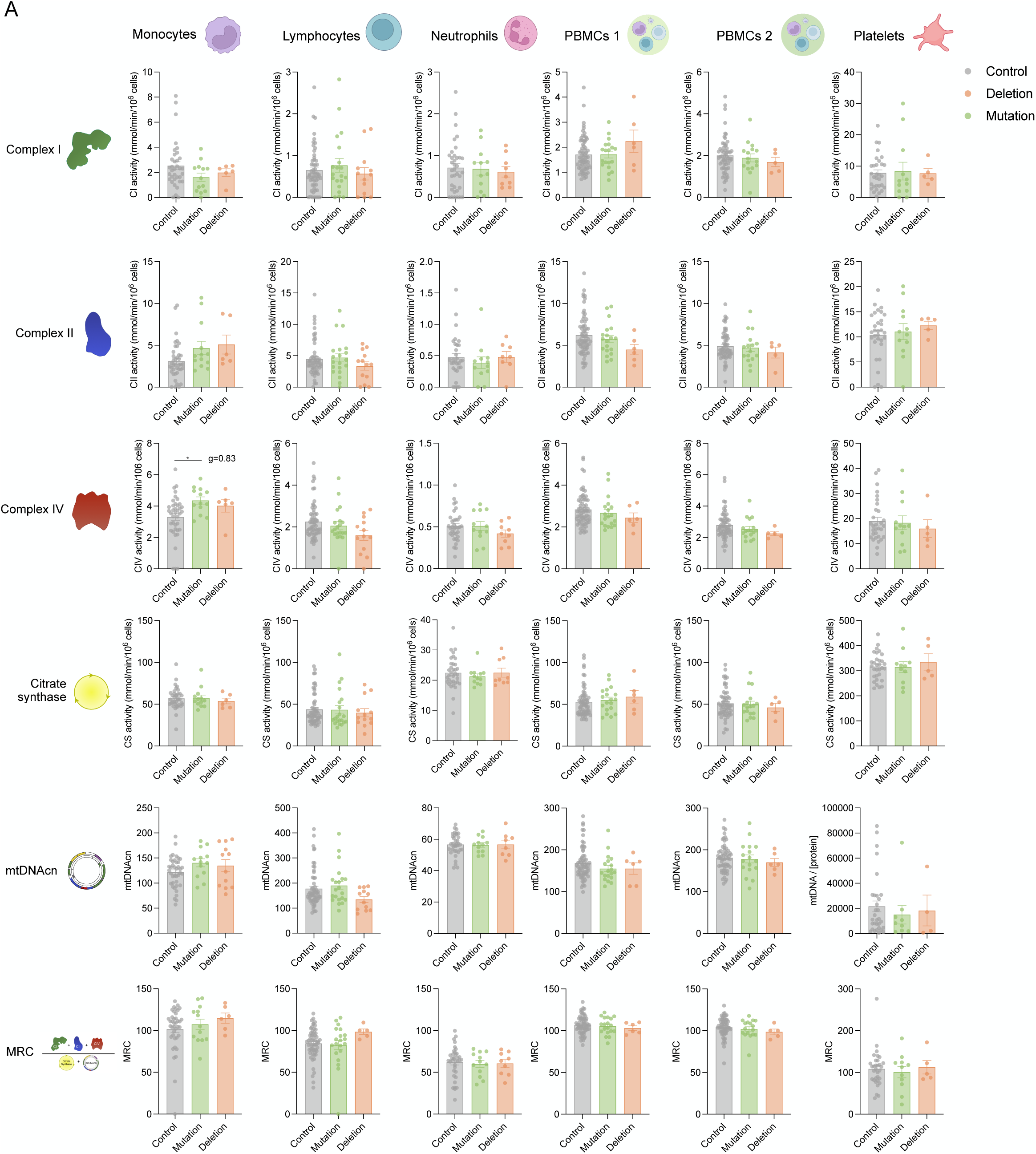
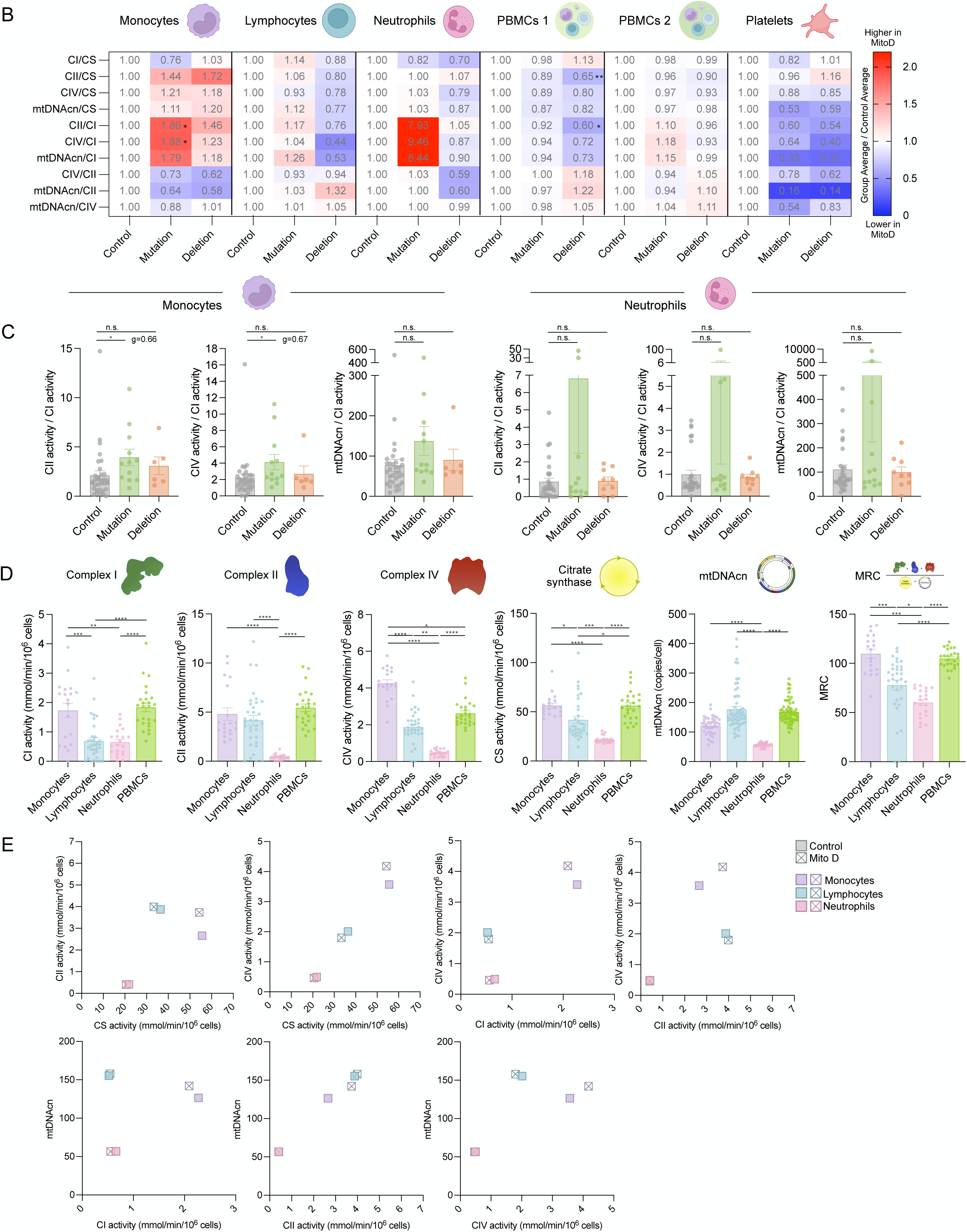
Differences in mitochondrial measures between healthy controls and MitoD groups per cell type. **(A)** Differences between mitochondrial enzyme activities, mtDNAcn, and MRC in Monocytes, Lymphocytes, Neutrophils, PBMCs and platelets between healthy controls and Mutation/MELAS and Deletion mitochondrial disease groups. Individual data points, mean +/- SEM shown. Effect size in Hedge’s g. **(B)** Heatmap of mean ratios of mitochondrial measures in MitoD groups relative to control in immune cell types, PBMCs and platelets, and **(C)** ratios of mitochondrial measures in MitoD groups relative to control in monocytes and neutrophils. **(D)** CI, CII, CIV, CS activity, mtDNAcn, MRC in immune cell types of mitochondrial disease patients. **(E)** Bivariate plots showing shift in the relationship between different pairs of mitochondrial enzyme measures and mtDNAcn in immune cell types between healthy controls and MitoD patients. Control n=31-66, Mutation n=11-21, Deletion n=5-13. One-way ANOVA, Kruskal Wallis multiple comparison test. *p<0.05, **p<0.01, ***p<0.001, ****p<0.0001.

**Figure S4.**
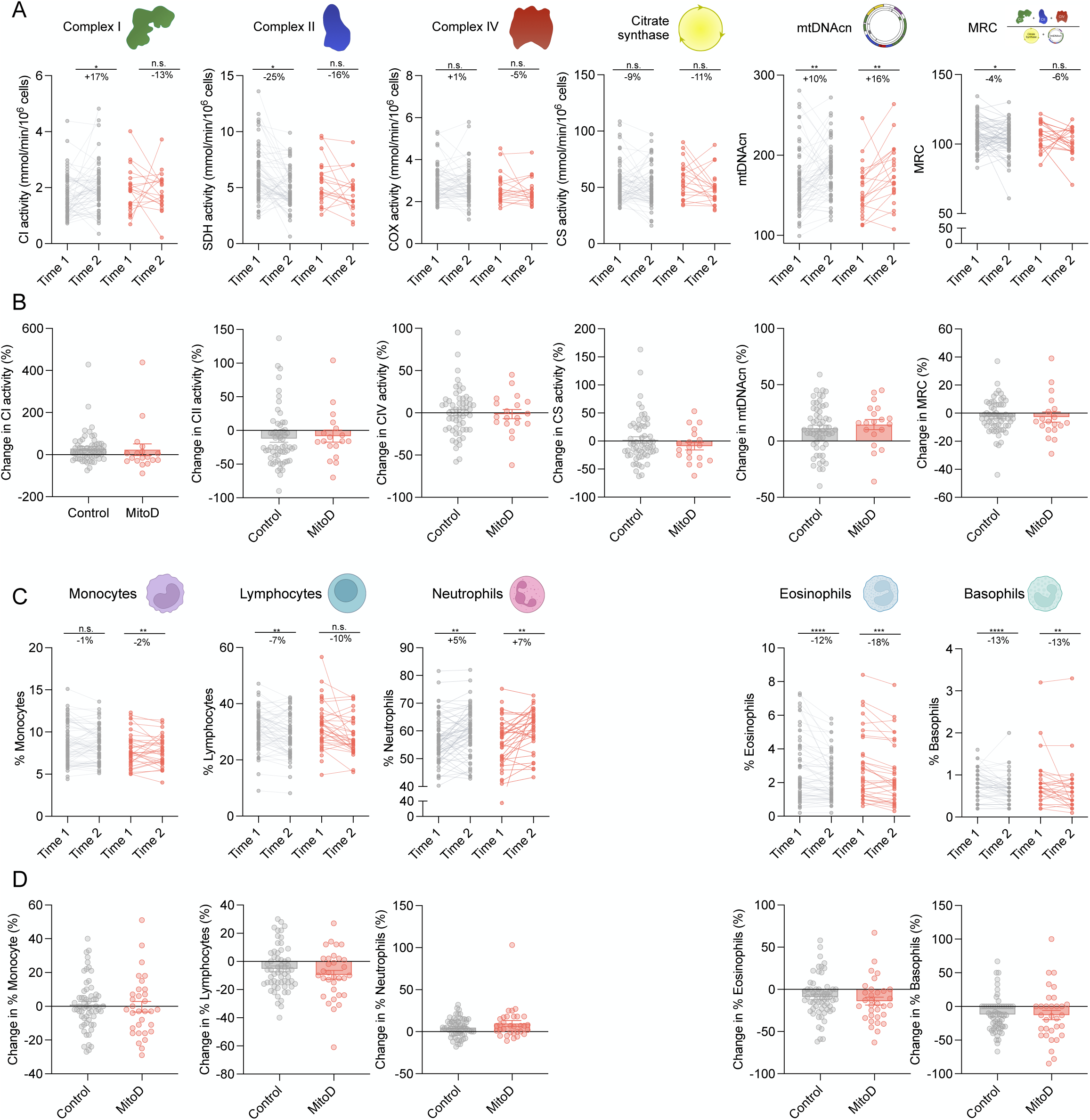
Shift in mitochondrial measures and WBC composition between different timepoints. Individual **(A)** trajectory of change and **(B)** percent changes in mitochondrial enzyme activities, mtDNAcn and MRC in healthy controls (grey) and MitoD patients (red). Individual **(C)** trajectory of change, with median % increase labelled, and **(D)** percent changes in % of WBCs of monocytes, lymphocytes neutrophils, eosinophils and basophils in healthy controls and MitoD patients, individual data points with mean % as bar. No significant differences were observed in any variable between MitoD and Control. Mitochondrial Features: Control n=58-59, MitoD n=18-19. WBC proportions: Control n=60, MitoD n=32. Wilcoxon test for paired timepoint measurements, Mann-Whitney test for group comparisons. *p<0.05, **p<0.01, ***p<0.001, ****p<0.0001. Individual data points, median +/- 95% confidence interval shown for comparison between groups.

**Figure S5.**
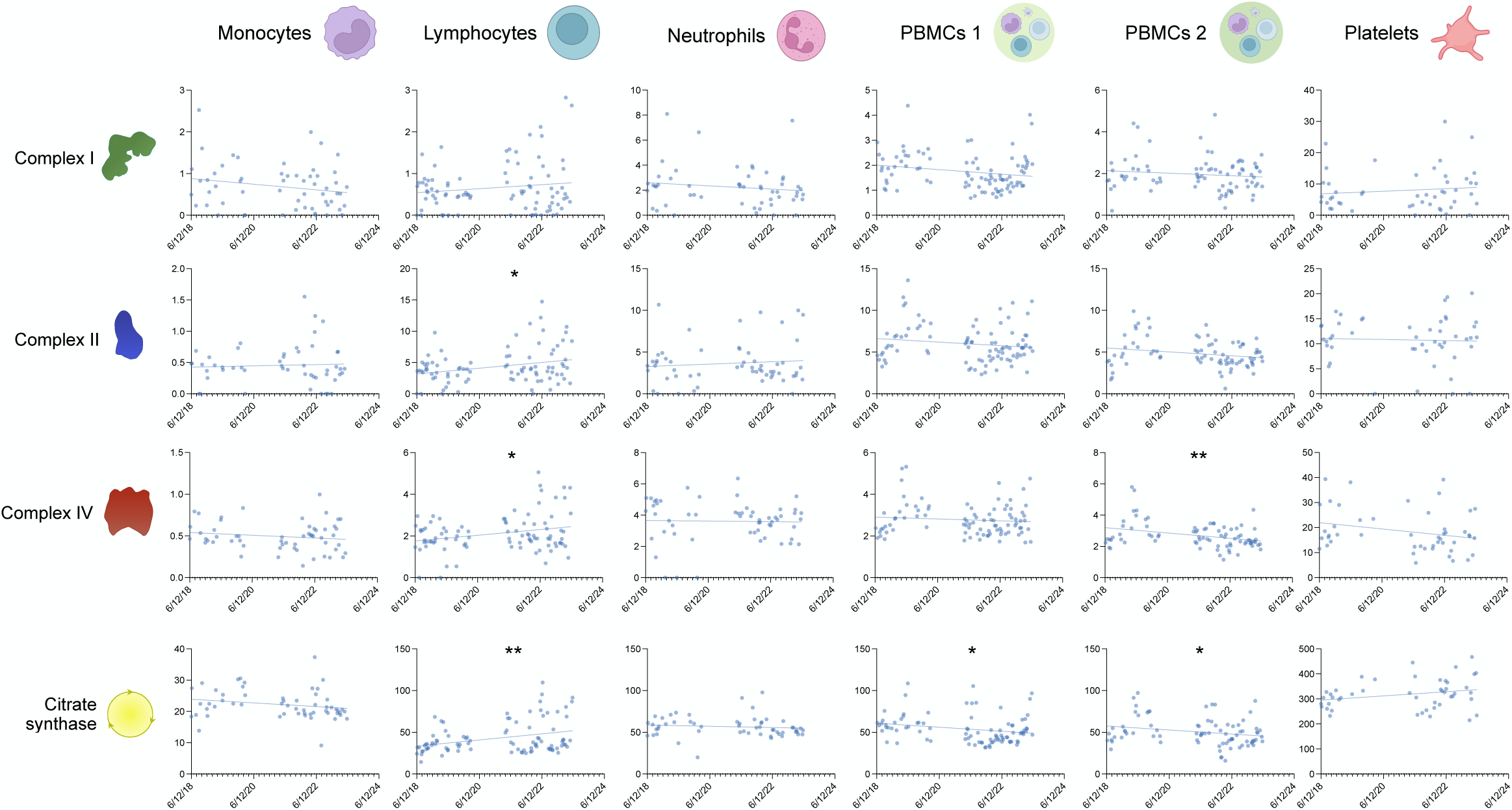
Effect of storage time on enzymatic activity assays. **(A)** CI, CII, CIV and CS activity of each cell type at different collection timepoints in monocytes, lymphocytes, neutrophils, PBMCs and platelets. Plots show enzyme activity on the y-axis, and draw date on the x-axis. Linear regression model line shown. n=47-92. Significant correlations between time and enzymatic activity are marked with asterisks. *p<0.05, **p<0.01, ***p<0.001, ****p<0.0001.

## REFERENCES

1. Jang JY, Blum A, Liu J, Finkel T. The role of mitochondria in aging. The Journal of Clinical Investigation 2018;128:3662–3670.

2. Picard M, Wallace DC, Burelle Y. The rise of mitochondria in medicine. Mitochondrion 2016;30:105–116.

3. Wilkinson MS, Dunham-Snary KJ. Blood-based bioenergetics: a liquid biopsy of mitochondrial dysfunction in disease. Trends in Endocrinology & Metabolism 2023;34:554–570.

4. Holper L, Ben-Shachar D, Mann JJ. Multivariate meta-analyses of mitochondrial complex I and IV in major depressive disorder, bipolar disorder, schizophrenia, Alzheimer disease, and Parkinson disease. Neuropsychopharmacology 2019;44:837–849.

5. Pyle A, Burn DJ, Gordon C, Swan C, Chinnery PF, Baudouin SV. Fall in circulating mononuclear cell mitochondrial DNA content in human sepsis. Intensive Care Medicine 2010;36:956–962.

6. Patti M-E, Corvera S. The Role of Mitochondria in the Pathogenesis of Type 2 Diabetes. Endocrine Reviews 2010;31:364–395.

7. Picard M, McEwen BS. Psychological Stress and Mitochondria: A Systematic Review. Psychosom Med 2018;80:141–153.

8. Tyrrell DJ, Bharadwaj MS, Jorgensen MJ, Register TC, Molina AJA. Blood cell respirometry is associated with skeletal and cardiac muscle bioenergetics: Implications for a minimally invasive biomarker of mitochondrial health. Redox Biology 2016;10:65–77.

9. Westerlund E, Marelsson SE, Karlsson M, et al. Correlation of mitochondrial respiration in platelets, peripheral blood mononuclear cells and muscle fibers. Heliyon 2024.

10. Devine J, Monzel AS, Shire D, et al. Brain-body mitochondrial distribution patterns lack coherence and point to tissue-specific and individualized regulatory mechanisms. bioRxiv 2024.

11. Nakano K, Tarashima M, Tachikawa E, et al. Platelet mitochondrial evaluation during cytochrome c and dichloroacetate treatments of MELAS. Mitochondrion 2005;5:426–433.

12. Ma YY, Li XY, Li ZQ, et al. Clinical, biochemical, and genetic analysis of the mitochondrial respiratory chain complex I deficiency. Medicine (Baltimore) 2018;97:e11606.

13. Ma YY, Wu TF, Liu YP, et al. A study of 133 Chinese children with mitochondrial respiratory chain complex I deficiency. Clinical Genetics 2015;87:179–184.

14. Ma YY, Wu TF, Liu YP, et al. Mitochondrial respiratory chain enzyme assay and DNA analysis in peripheral blood leukocytes for the etiological study of Chinese children with Leigh syndrome due to complex I deficiency. Mitochondrial DNA 2013;24:67–73.

15. Ma YY, Zhang XL, Wu TF, et al. Analysis of the mitochondrial complex I-V enzyme activities of peripheral leukocytes in oxidative phosphorylation disorders. J Child Neurol 2011;26:974–979.

16. Kotzailias N, Finsterer J, Zellner M, Marsik C, Dukic T, Jilma B. Platelet function in mitochondriopathy with stroke and stroke-like episodes. Thromb Haemost 2004;91:544–552.

17. Hubens WHG, Vallbona-Garcia A, de Coo IFM, et al. Blood biomarkers for assessment of mitochondrial dysfunction: An expert review. Mitochondrion 2022;62:187–204.

18. Walker MA, Lareau CA, Ludwig LS, et al. Purifying Selection against Pathogenic Mitochondrial DNA in Human T Cells. N Engl J Med 2020;383:1556–1563.

19. Walker MA, Li S, Livak KJ, Karaa A, Wu CJ, Mootha VK. T cell activation contributes to purifying selection against the MELAS-associated m.3243A>G pathogenic variant in blood. Journal of Inherited Metabolic Disease 2024;n/a.

20. Zhang J, Koolmeister C, Han J, et al. Antigen receptor stimulation induces purifying selection against pathogenic mitochondrial tRNA mutations. JCI Insight 2023;8.

21. Franklin IG, Milne P, Childs J, et al. T cell differentiation drives the negative selection of pathogenic mitochondrial DNA variants. Life Science Alliance 2023;6:e202302271.

22. Sanchez-Contreras M, Sweetwyne MT, Tsantilas KA, et al. The multi-tissue landscape of somatic mtDNA mutations indicates tissue-specific accumulation and removal in aging. eLife 2023;12:e83395.

23. Weiss SL, Henrickson SE, Lindell RB, et al. Influence of Immune Cell Subtypes on Mitochondrial Measurements in Peripheral Blood Mononuclear Cells From Children with Sepsis. Shock 2022;57.

24. Rausser S, Trumpff C, McGill MA, et al. Mitochondrial phenotypes in purified human immune cell subtypes and cell mixtures. eLife 2021;10:e70899.

25. Kramer PA, Ravi S, Chacko B, Johnson MS, Darley-Usmar VM. A review of the mitochondrial and glycolytic metabolism in human platelets and leukocytes: Implications for their use as bioenergetic biomarkers. Redox Biology 2014;2:206–210.

26. Chacko BK, Kramer PA, Ravi S, et al. Methods for defining distinct bioenergetic profiles in platelets, lymphocytes, monocytes, and neutrophils, and the oxidative burst from human blood. Laboratory Investigation 2013;93:690–700.

27. Maianski NA, Geissler J, Srinivasula SM, Alnemri ES, Roos D, Kuijpers TW. Functional characterization of mitochondria in neutrophils: a role restricted to apoptosis. Cell Death & Differentiation 2004;11:143–153.

28. Gordon-Lipkin EM, Banerjee P, Franco JLM, et al. Primary oxidative phosphorylation defects lead to perturbations in the human B cell repertoire. Frontiers in Immunology 2023;14.

29. Liu C-S, Cheng W-L, Lee C-F, et al. Alteration in the copy number of mitochondrial DNA in leukocytes of patients with mitochondrial encephalomyopathies. Acta Neurologica Scandinavica 2006;113:334–341.

30. Naue J, Xavier C, Hörer S, Parson W, Lutz-Bonengel S. Assessment of mitochondrial DNA copy number variation relative to nuclear DNA quantity between different tissues. Mitochondrion 2024;74:101823.

31. Wachsmuth M, Huebner A, Li M, Madea B, Stoneking M. Age-related and heteroplasmy-related variation in human mtDNA copy number. PLoS genetics 2016;12:e1005939.

32. Ackermann K, Revell VL, Lao O, Rombouts EJ, Skene DJ, Kayser M. Diurnal rhythms in blood cell populations and the effect of acute sleep deprivation in healthy young men. Sleep 2012;35:933–940.

33. Bushell D, Tan JKH, Smith J, Moro C. The identification of diurnal variations on circulating immune cells by finger prick blood sampling in small sample sizes: a pilot study. Laboratory Medicine 2023;55:220–226.

34. Lévi FA, Canon C, Touitou Y, et al. Circadian rhythms in circulating T lymphocyte subtypes and plasma testosterone, total and free cortisol in five healthy men. Clin Exp Immunol 1988;71:329–335.

35. Kelly C, Trumpff C, Acosta C, et al. A platform to map the mind–mitochondria connection and the hallmarks of psychobiology: the MiSBIE study. Trends in Endocrinology & Metabolism 2024;35:884–901.

36. Mosharov EV, Rosenberg AM, Monzel AS, et al. A Human Brain Map of Mitochondrial Respiratory Capacity and Diversity. bioRxiv 2024:2024.2003.2005.583623.

37. Picard M. Blood mitochondrial DNA copy number: what are we counting? Mitochondrion 2021;60:1–11.

38. Yatsuga S, Fujita Y, Ishii A, et al. Growth differentiation factor 15 as a useful biomarker for mitochondrial disorders. Ann Neurol 2015;78:814–823.

39. Suomalainen A, Elo JM, Pietiläinen KH, et al. FGF-21 as a biomarker for muscle-manifesting mitochondrial respiratory chain deficiencies: a diagnostic study. The Lancet Neurology 2011;10:806–818.

40. Warren C, McDonald D, Capaldi R, et al. Decoding mitochondrial heterogeneity in single muscle fibres by imaging mass cytometry. Scientific Reports 2020;10:15336.

41. Simard ML, Mourier A, Greaves LC, Taylor RW, Stewart JB. A novel histochemistry assay to assess and quantify focal cytochrome c oxidase deficiency. J Pathol 2018;245:311–323.

42. Picard M, Prather AA, Puterman E, et al. A Mitochondrial Health Index Sensitive to Mood and Caregiving Stress. Biol Psychiatry 2018;84:9–17.

43. Schaefer AM, Phoenix C, Elson JL, McFarland R, Chinnery PF, Turnbull DM. Mitochondrial disease in adults: A scale to monitor progression and treatment. Neurology 2006;66:1932–1934.

44. Sletten DM, Suarez GA, Low PA, Mandrekar J, Singer W. COMPASS 31: a refined and abbreviated Composite Autonomic Symptom Score. Mayo Clin Proc 2012;87:1196–1201.

45. Kaufmann P, Engelstad K, Wei Y, et al. Protean Phenotypic Features of the A3243G Mitochondrial DNA Mutation. Archives of Neurology 2009;66:85–91.

46. Beaudart C, Rolland Y, Cruz-Jentoft AJ, et al. Assessment of Muscle Function and Physical Performance in Daily Clinical Practice. Calcified Tissue International 2019;105:1–14.

47. Fisk JD, Ritvo PG, Ross L, Haase DA, Marrie TJ, Schlech WF. Measuring the functional impact of fatigue: initial validation of the fatigue impact scale. Clin Infect Dis 1994;18 Suppl 1:S79–83.

48. Walkowiak B, Kesy A, Michalec L. Microplate reader--a convenient tool in studies of blood coagulation. Thromb Res 1997;87:95–103.

